# The Sense of Body Ownership and the Neural Processes of Memory Encoding and Reinstatement

**DOI:** 10.1101/2023.03.07.531497

**Authors:** H. Iriye, M. Chancel, H. H. Ehrsson

**Affiliations:** Department of Neuroscience, Karolinska Institutet, Stockholm, Sweden; Université Grenoble Alpes, Université Savoie Mont Blanc, CNRS, LPNC, 38000, Grenoble, France

## Abstract

Every event we experience that results in a memory involves experiencing our conscious self at its heart. The spatial and perceptual experience of one’s own body is the most basic form of selfhood. Disrupting bodily selfhood during memory formation impairs functioning of the hippocampus during retrieval, which implies a weakening of encoding mechanisms. However, neural activity present as individuals encode memories during manipulations of bodily selfhood have yet to be assessed. We investigated how body ownership, a core aspect of bodily selfhood, impacts encoding processes in the hippocampus and additional core memory regions (e.g. posterior parietal cortex). Further, we assessed the degree to which memories are reinstated according to body ownership during encoding and vividness during retrieval as a measure of memory strength. Participants viewed immersive videos through VR glasses, which depicted everyday events that included a first-person view of a mannequin’s body aligned with participants’ real bodies during fMRI scanning. We manipulated feelings of body ownership over the mannequin using a perceptual full-body illusion. One week later, participants retrieved memories for the videos during fMRI scanning. We observed that patterns of activity in several regions including the hippocampus and posterior parietal cortex distinguished between videos encoded with strong versus weak illusory body ownership. Moreover, these same regions reinstated memories to a greater degree when they were formed with a strong sense of body ownership and retrieved with increasing levels of vividness. Our findings demonstrate how the fundamental sense of owning one’s body shape neural signatures of memory formation and strengthen reinstatement at retrieval.

---

Every time we experience or remember an event, our conscious self is at the centre of the episode (Tulving, 1985). This intimate association between the conscious self and memory suggests they are fundamentally related. The spatial and perceptual experience of the body as one’s own (i.e. body ownership) is a basic form of conscious self-experience (Blanke et al 2015), and rooted in a coherent multisensory representation of one’s body based on the continuous integration of bodily-related sensory signals, including touch, vision, proprioception, and interoception (Ehrsson, 2020). This perceptual bodily self defines the egocentric reference frame that is crucial for the processing of sensory and cognitive information. Emerging literature suggests that the multisensory experience of one’s own body at encoding influences memory upon retrieval (Bergouignan et al., 2014; Bréchet et al., 2019, 2020; Iriye & Ehrsson, 2022; Tacikowski et al., 2020). The idea is that the perception of one’s own body binds sensory and cognitive information into a unified experience with the self in the centre during encoding, and that impairing this fundamental binding weakens the effectiveness of the encoding process leading to reduced quality of memories at recall. Behavioural studies that manipulated the sense of body ownership by using bodily illusions during encoding (Iriye & Ehrsson 2021; Tacikowski et al., 2020) found reductions in vividness and memory accuracy during later recall for events that were encoded with a reduced sense of body ownership. Bergouignan and colleagues (2014) observed a reversal of the normal activation pattern of the left posterior hippocampus during repeated retrieval of social interactions encoded while experiencing an out-of-body illusion, which correlated with reduced vividness. However, hippocampal activity was not measured at encoding and the specific effect of body ownership cannot be disentangled from effects of changes in self-location and visual perspective that are also components of the out-of-body illusion (Ehrsson, 2007; Guterstam & Ehrsson, 2012). Furthermore, it has been theorized that the angular gyrus in the ventral posterior parietal cortex may integrate body ownership within memory (Bréchet et al., 2018). The angular gyrus supports multisensory encoding (Bonnici et al., 2016; Jablonowski & Rose, 2022) by combining cross-modal information into a common egocentric framework (Bonnici et al., 2018; Humphreys et al., 2021), and encodes self-relevant stimuli (Singh-Curry & Husain, 2009). However, whether angular gyrus activity reflects body ownership during memory encoding has not been directly tested.

During retrieval, there is a reinstatement of brain activity patterns that were present when events were initially experienced, which is coordinated by the hippocampus (e.g. Hebscher et al., 2021). The higher the overlap between patterns of neural activity at encoding and retrieval, the stronger the memory trace (Bird et al., 2015; Oedekoven et al., 2017). Hence, if the hippocampus is sensitive to own-body perception during encoding, as has been hypothesised (Bergouignan et al., 2014), this may have consequences for the strength of memory reinstatement in a manner that scales with memory vividness. Experiencing naturalistic multisensory memories with a strong sense of body ownership during encoding could facilitate access to hippocampal representations of the memory trace during vivid recall, which would lead to increased reinstatement of these memories in brain regions initially involved in encoding across the cortex.

Here we investigate how changes in body ownership during encoding infuence neural processes related to memory encoding and memory reinstatement. Participants watched immersive stereoscopic videos of naturalistic scenes through a head-mounted display (HMD) during functional magnetic resonance imaging (fMRI) scanning while we manipulated their feelings of ownership over a mannequin’s body in the centre of the scene with synchronous/asynchronous visuotactile stimulation in a well-established full-body illusion paradigm (Petkova et al 2008; Ehrsson 2020). One week later, participants recalled these experiences during functional scanning. We predicted that patterns of activity in the hippocampus and angular gyrus during encoding would reflect whether an event was experienced with a strong or weak sense of body ownership of the artificial body in the scene. We also predicted that memory reinstatement in the hippocampus and the distributed cortical regions associated with the multisensory experience during encoding, would be greater for memories formed with illusory body ownership and high vividness during retrieval.

## Methods

### Participants

Participants included 30 healthy, right-handed young adults (age range 19-29 years old) with no prior history of neurological or psychiatric impairment, and who were not currently taking medication that affected mood or cognitive function (18 men, 12 women; mean age = 25.13, *SD* = 3.06; mean years of education: 16.31, *SD* = 3.01). Importantly, we only recruited participants who had not previously taken part in any experiments involving body illusions to ensure they would be naïve to the illusion. Participants provided informed written consent as approved by the Swedish Ethical Review Authority. Six participants were excluded from the fMRI analyses due to technical issues during scanning (i.e., scanner overheating, N = 1) and excessive head movement during the memory encoding session (i.e. > 3mm in pitch, yaw, and/or roll, N = 5). Thus, the final fMRI analyses was performed on 24 participants (15 men, 9 women; mean age = 25.33 years old, *SD* = 3.31; mean years of education = 16.31, *SD* = 3.14). The sample size in the present study (*N* = 24) is comparable to the sample size of Bergouignan and colleagues (2014; *N* = 21).

### Materials

Stimuli consisted of 26 high-resolution digital 3D-videos depicting everyday life events in various famous locations around Stockholm, Sweden that have been described in a previous experiment (Iriye & Ehrsson, 2022). We chose recognizable local places of interest to recreate natural real-life experiences for participants and enhance the ecological validity of our study. Each video included a stereoscopic view of a mannequin’s body seen from a supine, first-person perspective. Stereoscopic vision was achieved by filming each event with two GoPro Hero 7 cameras (San Mateo, California, USA) mounted side-by-side on a tripod to mimic left and right eye positions. The recordings from the left and right cameras were viewed in the left and right eye of a pair of MR-compatible LCD displays in the HMD (VisualSystem HD, Nordic Neurolab, Norway) to create a stereoscopic effect within the videos. Throughout each video, a white Styrofoam ball (6.5 cm in diameter) attached to a wooden stick (1 m long) repeatedly stroked the mannequin’s torso from the sternum to the belly button in a downwards direction every 2 seconds for 40 seconds total (Iriye & Ehrsson, 2022; O’Kane and Ehrsson, 2021). Each stroke lasted approximately one second and lasted the duration of each video. After 20 seconds, an everyday scene unfolded in the video, which always involved the same two actors engaged in a short conversation based on a unique, visible item. One of the actors was an author of the present study referred to as “Heather” in the videos, and the second was a fellow lab member referred to as “Vicki”. The background of each scene consisted of a typical summer day in a well-known location in Stockholm, Sweden (e.g. Central Station, Gamla Stan, Kungsträdgården) complete with members of the public going about their lives as usual. For example, in the video titled “Central Station”, Vicki and Heather are walking toward the main entrance of the city’s main train station together discussing a vacation they are about to take to Oslo, Norway. The two discuss details of the trip including how long they’ll be gone for and who they are staying with. Vicki pauses and mentions she thinks she has left her travel pillow at home. She removes the backpack she is wearing, rifles through it to check, finds she has packed it after all, and shows a blue travel pillow to Heather. Then the two continue to make their way to the entrance of the station. A grey van is parked on the street to the right and two green bicycles are locked to poles next to the station. Several people are seen entering and exiting the station in the background. The video footage of the mannequin being stroked by the wooden stick was filmed separately against a green screen and imposed onto the footage of the everyday scenes using Final Cut Pro X 10.4.7. An audio track of the dialog was recorded separately from the video in a soundproof environment using a Røde Videomic Go microphone (Sydney, Australia). Background noises appropriate to each video (e.g., birds, wind, distant conversations, traffic) were downloaded from an open source repository (https://pixabay.com/sound-effects/search/background) and layered underneath the dialog using Audicity ® 2.3.2. The final audio track was integrated with its respective video using Final Cut Pro X 10.4.7. A second audio track only heard by the experimenter through a separate set of headphones was added to each video to cue precise timing and duration of the tactile stimuli. Audio tracks delivered to the experimenter and participant were separated by a Edirol USB AudioCapture sound card (model number: UA-25EX, 24-bit, 96 kHz) and played through MR-compatible headphones (AudioSystem, Nordic Neurolab, Norway).

We adopted the same procedure to ensure that any observed behavioural or fMRI effects of experimental condition would not be due to differences in the complexity of the individual videos as described in our previous study (Iriye & Ehrsson, 2022). Two independent raters judged each video in terms of multiple sub-categories of complexity and composite scores of each sub-category were used to divide videos into two groups (see Supplemental Table 1) to minimize differences in average complexity between groups (see Supplemental Table 2; ; Bonasia, Sekeres, Gilboa, Grady, Winocur, and Moscovitch, 2018; Iriye and Ehrsson, 2022; Sekeres, Bonasia, St-Laurent, Pishdadian, Winocur, Grady, and Moscovitch, 2016). Assignment of each video to an experimental condition was counterbalanced across participants. The individual raters judged each video in terms of its visual, audio, narrative, and emotional complexity, using five-point Likert scales (One = Low, Five = High). Visual complexity was measured along three dimensions concerning complexity of the background scene, amount of movement, and number of characters. Auditory complexity referred to the complexity of the background audio, while narrative complexity referred to complexity of the storyline. Emotional complexity was characterized according to degree of sadness, excitement, joy, anger, disgust, fear, and shame present within each video. A two-way random effects model was run for each complexity dimension to assess inter-rater reliability. The emotional sub-categories of sadness, anger, disgust, fear, and shame did not display enough variance to run the analysis as all ratings from both raters were at floor level (i.e., one). The average intraclass correlation coefficient across complexity categories was 0.78 (*SD* = .08), demonstrating a reliable degree of agreement between raters (see Supplemental Table 3 for means from each sub-category).

### Procedure

#### Session One: Memory Encoding

The study involved two separate fMRI scanning sessions spaced seven days apart. During the first session, participants were fitted with a HMD containing a pair of MR-compatible LCD visual displays (VisualSystem HD, Nordic Neurolab, Norway) and repeatedly watched the 24 of the immersive videos during functional scanning. The location of the participants’ real bodies lying on the scanner table was aligned with a first-person view of the mannequin’s body in the video (i.e., the experimenter ensured that the participants’ arms and legs were straight and the participants’ body was located on the centre of the table to match the mannequin in the videos). Each video was associated with a unique title (i.e. the location depicted in the scene) that was presented in the centre of the screen for 2.5 seconds before the video began. To induce a bodily illusion of the mannequin feeling like one’s own body with propriopceptive and tactile sensations seemingly originating from the fake body, participants were instructed to relax and focus on the mannequin as touches were delivered on their torso at the same time and in the same direction that they saw the mannequin’s torso being touched in the video, i.e., synchronous visuotactile stimulation (e.g. Petkova & Ehrsson, 2008). As a control condition, participants saw and felt touches in an alternating temporal pattern in the other half of the videos, i.e., asynchronous visuotactile stimulation, which significantly reduces the illusion (Petkova & Ehrsson, 2008). Critically, the strokes in both conditions were identical in magnitude and location, and the only difference between conditions was the one second delay between seen and felt strokes in the asynchronous condition.

Participants were instructed to remember as much as possible about each video, including the title. Videos were separated by an active baseline consisting of a left versus right decision task that lasted between 2.5–10 seconds and was distributed such that shorter inter-trial intervals occurred more frequently than longer inter-trial intervals (4 x 2.5s, 3 x 5s, 2 x 7s, 1 x 10s; Iriye & St. Jacques, 2020; Stark & Squire, 2001). We chose an active baseline as activity in the medial temporal lobes can be higher when participants rest in between trials compared to when they are performing a memory task during a trial, which can be reversed by adopting a left-right decision task as a baseline measure (Stark & Squire, 2001). During rest, participants have the opportunity for mind-wandering and self-reflection that can activate core memory regions. Thus, having participants perform a mindless task such as deciding whether an arrow is pointing left or right is a more appropriate baseline task to compare against memory encoding and retrieval.

Participants viewed individual videos a total of three times over the course of nine functional runs, each consisting of eight videos, for a total of 36 trials per condition. Videos were repeated three times to ensure that participants would be able to recall each video one week later during session two, which had been determined by pilot testing on a separate group of participants (data not shown). Video presentation was pseudo-randomized across runs, such that no condition was repeated more than twice in a row and videos repeated every three runs. Prior to scanning, the experimenter verified that the participant understood the task (i.e. to remember as much as possible about each video, including the title, dialogue, and visual scene) by having the participant verbally repeat the instructions. Participants were also immersed within two videos not used for the main memory experiment, one with synchronous visuotactile stimulation and one with asynchronous visuotactile stimulation, inside a mock scanner to become accustomed to the experimental stimuli and active baseline task.

#### Session One: Cued Recall and Subjective Ratings

Immediately after scanning, participants completed a cued recall test comprised of five questions per video to assess objective memory accuracy outside the scanning environment in a nearby testing room. Three questions were related to the main storyline (e.g. Which animal in the museum was Heather surprised to see?) and two questions concerned background peripheral details not important to the storyline (e.g. How many scooters were parked outside the museum?). To assess memory phenomenology, participants rated reliving, emotional intensity, vividness, and degree of belief in memory accuracy on a seven-point Likert scales (1 = None, 7 = High; Iriye & Ehrsson, 2022; Iriye & St. Jacques, 2021). Cued recall questions and subjective ratings were completed on a computer. Cued recall questions from each video were randomly assigned to either session one or session two for each participant and presented in random order.

#### Session Two: Memory Retrieval

One week after session one, participants were asked to repeatedly retrieve each of the videos from memory. We chose a one-week interval between testing sessions to maintain consistency with previous studies in the field, which have shown declines in the richness of recollection during retrieval over a delay of this length (Bergouignan et al., 2014; Iriye & Ehrsson, 2022). Each trial began with the video title presented in the centre of the field of view for 2.5s, followed by an instruction for participants to close their eyes and retrieve the memory in as much detail as possible for a duration of 17.5s. After 17.5s had elapsed, an auditory cue sounded through the headphones which signalled participants to stop retrieving their memory and open their eyes (2.5s). Immediately following each trial, participants were asked to rate the vividness of their memory on a visual analogue scale (0 = Low, 100 = High). Participants had 2.5s for the vividness rating and responded using an MR-compatible mouse. Each video was retrieved a total of three times over the course of six functional runs (12 trials each), resulting in a total of 36 trials per condition. Trials were separated by the same active baseline task as session one, adapted to the increased number of trials per functional run (6 x 2.5s, 3 x 5s, 2 x 7.5s, 1 x 10s). Video presentation was pseudo-randomized such that a given condition was not repeated more than twice in a row.

Prior to scanning, participants were presented with the video titles and asked to indicate whether they were able to recall the associated video with a yes/no response to verify that they would be successfully retrieving a memory during the experiment. In the event that the response was “no”, the participant received a word cue related to the theme of the video (e.g. “picnic” for the video Hagaparken) and was asked to indicate if this helped them recover a specific memory. If the response was still “no”, the participant was instructed to focus on trying to recover their memory for that video during the 17.5 s retrieval period in the scanner. Participants were unable to retrieve a memory for 23 of the 576 total videos (.04% of total trials). The experimenter also verified that the participant understood the task (i.e. to close their eyes and vividly recall the video in as much detail as possible – not just the words that were said, but details of the surrounding, objects, and people in the scene) by having the participant verbally repeat the instructions before entering the scanner.

#### Session Two: Illusion Testing

We measured the strength of the full body illusion during session two, directly after participants had finished retrieving their memories of the videos. Participants were immersed within two previously unseen videos that were not viewed during the main memory portion of the experiment, one with synchronous and the other with asynchronous visuotactile stimulation, while lying on the MR scanner bed to mimic how the videos presented during the encoding session were experienced. However, the scanner was not recording functional images during illusion testing. Video presentation order was counterbalanced across participants. Each video was viewed twice. These new videos were shown purely to confirm that the bodily illusion manipulation worked as expected and that a stronger illusion was experienced in the synchronous compared to asynchronous condition. The videos were similar to those used in the main experiment: they involved an initial 20 seconds of a mannequin’s body viewed from a first-person perspective being stroked on the torso within a static scene depicting a unique location around Stockholm, Sweden, followed by an additional 20s of two characters engaged in an everyday conversation. However, unlike the videos included in the main experiment a knife appeared and slid quickly just above the mannequin’s stomach, which lasted approximately 2 seconds, at two points in each video (i.e. at 18s and 32s). There was no tactile stimulation applied to the participants’ bodies during the knife threat or for two seconds following it. The knife threats trigger stronger physiological emotional defence reactions that translate to increased sweating that can be registered as changes in skin conductance when the illusion is experienced compared to when it is not. This threat-evoked skin conductance (SCR) response serves as an indirect psychophysiological index of the illusion (Petkova et al., 2008; Ehrsson et al., 2007; Guterstam et al., 2015; Tacikowski et al., 2020). We did not include knife threats in the videos used for the main experiment to avoid confounding effects on memory.

We recorded threat-evoked skin conductance responses (SCR) with the MR-compatible Biopac System (MP160, Goleta, CA, USA; sampling rate = 100 Hz). The data were processed with AcqKnowledge® software (Version 5.0, Biopac). Two electrodes with electrode gel (Biopac, Goleta, CA, USA) were placed on the participants’ left index and middle fingers (distal phalanges). The script used to present the videos created in PyschoPy (v3.2.4; Peirce et al., 2019) sent a signal to the recording software to mark the beginning of each knife threat. We also tracked participants’ eye movements while they watched the videos used to assess the strength of the illusion induction to determine whether there were any systematic differences in gaze patterns between the synchronous and asynchronous conditions that may potentially have influenced the results of the memory task. MR-compatible binocular eye tracking cameras mounted inside of the VisualSystem HD (acquisition frequency: 60 Hz; NordicNeuroLab, 2018, Norway) recorded fixation co-ordinates as participants watched the videos.

Immediately after participants had viewed the first illusion testing video, they were led to the MR control room and completed two sets of questionnaires that assessed the degree of illusory bodily ownership they felt over the mannequin and the level of presence they felt within the immersive scene presented in the HMD (see further below). The illusion induction questionnaire consisted of three statements pertaining to the degree of multisensory bodily illusion of the mannequin as one’s own body and three control statements on seven-point Likert scales (−3 = Strongly Disagree, 0 = Neutral, 3 = Strongly Agree; see Table 1) presented in random order on a computer screen. S1 and S2 assessed “referral of touch” from the participant’s actual body to the mannequin indicating coherent multisensory binding of the seen and felt strokes on the mannequin’s body, while S3 assessed degree of explicitly sensed ownership over the mannequin’s body (Petkova & Ehrsson, 2008). The presence questionnaire was included to monitor the overall feeling that participants experienced “being there” inside the immersive 3D scene, which we reasoned might have unintended effects on participants’ ability to recall specific details about each scene, independent from the bodily illusion illusion. The presence questionnaire was comprised of three statements that participants rated on seven -point Likert Scales adapted from Slater (2000; see Table 2). The same process was repeated for the second illusion testing video associated with the opposite condition (i.e. if the first illusion testing video involved synchronous visuotactile stimulation, the second illusion testing video involved asynchronous visuotacitle stimulation). After the participants had completed the illusion testing and questionnaires for the second illusion testing video, they returned to the scanner bed a final time, where they watched the same two videos back-to-back in the same order as we measured their skin conductance responses and eye movements (as described above). Thus, for each condition we obtained one set of questionnaire ratings, skin conductance data from four knife threats (i.e. two per video), and eye movement data from both presentations of each video. We counterbalanced the order of video presentation and the condition assigned to each video across participants.

**Table 1.** Illusion Testing Questionnaire

|  |  |
| --- | --- |
| **S1** | I felt the touch of the brush on the mannequin in the location where I saw the brush moving |
| **S2** | I experienced that the touch I felt was caused by the brush touching the mannequin |
| **S3** | I felt as if the mannequin I saw was my body |
| **S4** | I could no longer feel my body |
| **S5** | I felt as if I had two bodies |
| **S6** | When I saw the brush moving, I experienced the touch on my back |

**Table 2.**
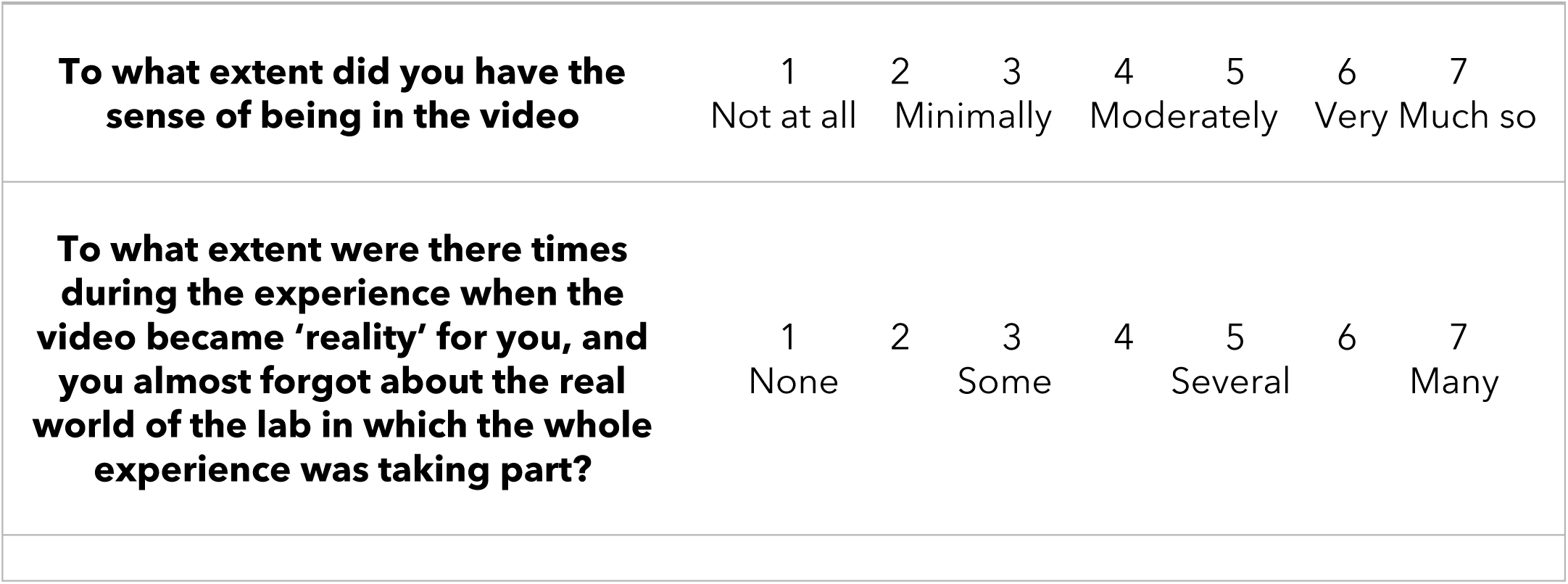
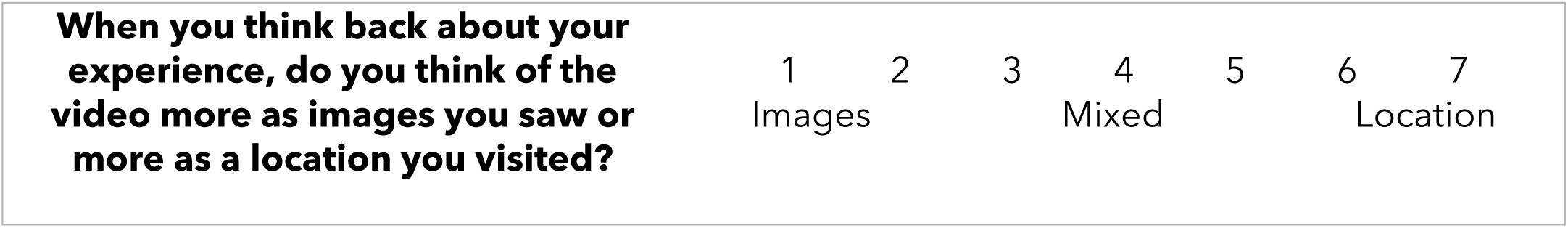
Presence Questionnaire

#### Session Two: Cued Recall and Subjective Ratings

At the end of session two, participants completed a cued recall accuracy test with new questions and the same subjective ratings made during session one outside of the scanning environment in a nearby testing room on a laptop. Cued recall questions were randomly assigned to either session one or session two for each participant and presented in random order. The instructions and procedure for completing the cued recall and subjective ratings were identical to those for session one.

#### MRI Data Acquisition and Preprocessing

Functional and structural images were collected on a GE750 3 Tesla MRI scanner. Detailed anatomical data were collected using a T1-weighted BRAVO sequence. Functional images were acquired using at T2*-weighted echo planar sequence (TR = 2500ms, TE = 30ms, FOV = 230 x 230, slice thickness = 3 mm). Whole brain coverage was obtained via 48 coronal oblique slices, acquired at an angle corresponding to AC-PC alignment in an interleaved ascending fashion, with a 2.4 x 2.4 mm in-plane resolution. The first ten volumes of each run were discarded to allow for T1 equilibrium.

Preprocessing of functional images was performed using SPM12 (Wellcome Department of Imaging Neuroscience, London, UK). Functional images were corrected for differences in acquisition time between slices for each whole brain volume using slice-timing, realigned within and across runs to correct for head movement, spatially normalized to the Montreal Neurological Institute (MNI) template (resampled at 2 x 2 x 2 mm voxels), and spatially smoothed using a Gaussian kernel (3mm full-width at half maximum). BOLD signal response patterns according to condition for both the encoding and retrieval scanning session separately were estimated using general linear models (GLMs). A regressor was estimated for every trial in each functional run to facilitate the multivariate analyses (see below), resulting in eight beta estimates (i.e. four per condition) for each of the nine functional runs in the encoding session and twelve beta estimates (i.e. six per condition) for each of the six functional runs in the retrieval session. To provide the multivariate classifiers with as much training data as possible for each condition and improve classification accuracy, we calculated response estimates for each trial (Turner, Mumford, Poldrack, & Ashby, 2012), as opposed to one mean estimate for each condition in a run as is standard for GLM analyses. Thus, there were a total of 36 beta estimates per condition for each scanning session. For the encoding session, regressors were time-locked to the onset of the natural social event in the video signalling the beginning of the memory task, after the initial 20s of visuotactile stimulation that was only used to trigger the bodily illusion (and no memorable event was presented). The duration lasted until the end of the video (i.e. 20s). We excluded the initial 20s of each video from the main analyses of interest to allow time to induce (i.e. synchronous condition) illusory ownership of the mannequin’s body, which has been previously shown to occur approximately within this time period (Petkova & Ehrsson, 2011;O’Kane et al 2021). The corresponding initial period of 20s visuotactile stimulation was also from the asynchronous condition to match the conditions in this aspect. For the retrieval session, regressors were time-locked to the onset of the memory retrieval cue and the duration was set to cover the retrieval period (i.e. 17.5s), excluding the auditory tone and the vividness rating. For both sessions, six movement parameters were included as separate regressors. The SPM canonical haemodynamic response function was used to estimate brain responses. Trial-wise beta estimates were used as input for the decoding analyses conducted on the encoding and retrieval sessions. For the representational similarity analysis (RSA), we averaged beta estimates across the three repetitions of each video (*N* = 24) for each scanning session separately to allow us to compare patterns of activity at encoding and retrieval according to condition i.e., illusion (synchronous) or control (asynchronous).

### Behavioural Data Analysis

To assess the strength of the bodily illusion induction from the questionnaire data, we calculated illusion statement ratings corrected for unspecific cognitive effects (e.g. confabulation and suggestibility) by subtracting average control statements from average illusion statement ratings separately for each condition (Iriye & Ehrsson, 2021). To analyze skin conductance responses collected during illusion induction testing, we used a custom Matlab toolbox (Tacikowski, Weijs, & Ehrsson, 2020) to identify the amplitude of each response defined as the difference between the maximum and minimum conductance values 0-6 s after a knife threat (Tacikowski, Weijs, & Ehrsson, 2020). We then performed a paired-samples t-test comparing magnitudes of skin conductance responses to knife threats between conditions. All trials were included in the skin conductance response analysis, i.e., also null responses, which means that we assessed the magnitude of the SCR for each condition (Dawson, 2000). To determine whether or not there were differences in gaze behaviour and overt attention due to experimental condition, we analyzed eye tracking data by calculating average fixation co-ordinates for both conditions for each participant, and then conducting a 2 (visuotactile stimulation: synchronous, asynchronous) x 2 (fixation coordinate: X, Y) repeated-measures ANOVA (Guterstam et al., 2015). We were unable to collect SCR data from four participants and eye tracking data from two participants due to technical issues with the recording equipment. Thus, the analysis of SCR data was conducted on 26 participants and the analysis of the eye tracking data was conducted on 28 participants.

Cued recall questions were coded for accuracy using strict criteria in which responses had to exactly match the correct response to be scored as correct (e.g. *What type of injury was Heather recovering from?* Correct answer: *A sprained ankle*, Incorrect answer: *A foot injury*). Using a more lenient criteria that allowed for partial marks did not change the overall pattern of results. The percentage of correct responses for central and peripheral details for both conditions and testing points was calculated for each participant. Subjective ratings of memory phenomenology were averaged across videos to create a mean score for each rating of reliving, emotional intensity, vividness and degree of belief in memory accuracy (see above).

Shapiro-Wilk tests were used to assess normality of the data. We used repeated measures ANOVAs to analyze data that was normally distributed and Wilcoxon signed-rank tests to assess data that was not normally distributed. Follow up t-tests were corrected for multiple comparisons using Bonferroni corrections. Greenhouse Geisser corrections were applied where the data violated assumptions of sphericity. Significance was defined as *p* < .05. Statistical analyses of all behavioral data were conducted using JASP (2019).

### fMRI Analyses

#### Multivariate Pattern Analysis (MVPA) of Encoding and Retrieval Sessions

All decoding analyses were performed with the CoSMoMVPA toolbox (Oosterhof et al., 2016). For each scanning session, we conducted a wholebrain searchlight analysis to identify regions where patterns of neural activity could distinguish between the synchronous (illusion) and asynchronous (control) conditions during the memory encoding and retrieval sessions separately based on trial-wise beta estimates. For each subject, a sphere comprised of 100 voxels was fitted around each voxel in the acquired volumes to create searchlight maps for each participant that reflected classification accuracies determined by k-fold cross-validation using a linear discriminant analysis (LDA) classifier. We investigated group-level effects corrected for multiple comparisons by submitting the resulting subject-specific classification accuracy maps to threshold-free cluster enhancement (TFCE). TFCE calculates a combined value for each voxel after a raw statistical map has been thresholded over an extensive set of values (Smith & Nichols, 2009). This approach capitalizes on the heightened sensitivity of cluster-based thresholding methods, without the limitation of defining an initial fixed cluster threshold a-priori (Oosterhoff, Connolly, & Haxby, 2016; Smith & Nichols, 2009), and is less affected by non-stationarity within data (Salimi-Khorshidi, Smith, & Nichols, 2011). 10,000 Monte Carlo simulations were estimated to identify brain regions where classification accuracy was higher than chance level (i.e. 50%). The resulting group-level wholebrain classification accuracy map was thresholded at *z* = 2.05 to correspond to a false discovery rate of *q* < .05 (q is the adjusted p-value after using the FDR approach). Significant clusters of activation were extracted and anatomically labelled using the xjview toolbox (https://www.alivelearn.net/xjview). The anatomical labels were then cross-referenced with the Yale Bioimage Suite MNI <-> Talaraich Tool (https://bioimagesuiteweb.github.io/webapp/mni2tal.html) for accuracy and manually compared to the mean normal structural scans of the current group of participants.

We then performed two separate region of interest (ROI) analyses (i.e., one for each scanning session) to investigate whether patterns of activity in the hippocampus could predict whether a memory was associated with the body ownership illusion (synchronous) or not (asynchronous) during memory encoding and retrieval. The hippocampal ROI was created by forming a 10mm sphere centred on co-ordinates reported by Bergouignan and colleagues (2014; X= -27, Y = -31, Z = -11; see Figure 1) using the MarsBar toolbox for SPM (Brett et al., 2002). The ROI was converted to binary format and resampled to 2mm cubic voxels. Decoding accuracy for each participant and for each session (i.e. encoding, retrieval) within the hippocampal ROI was estimated using a LDA classifier. Subject-level accuracies were then submitted to a two-sided, one-sample t-test against a null hypothesis of chance-level classification accuracy (i.e. 50%).

**Figure 1.**
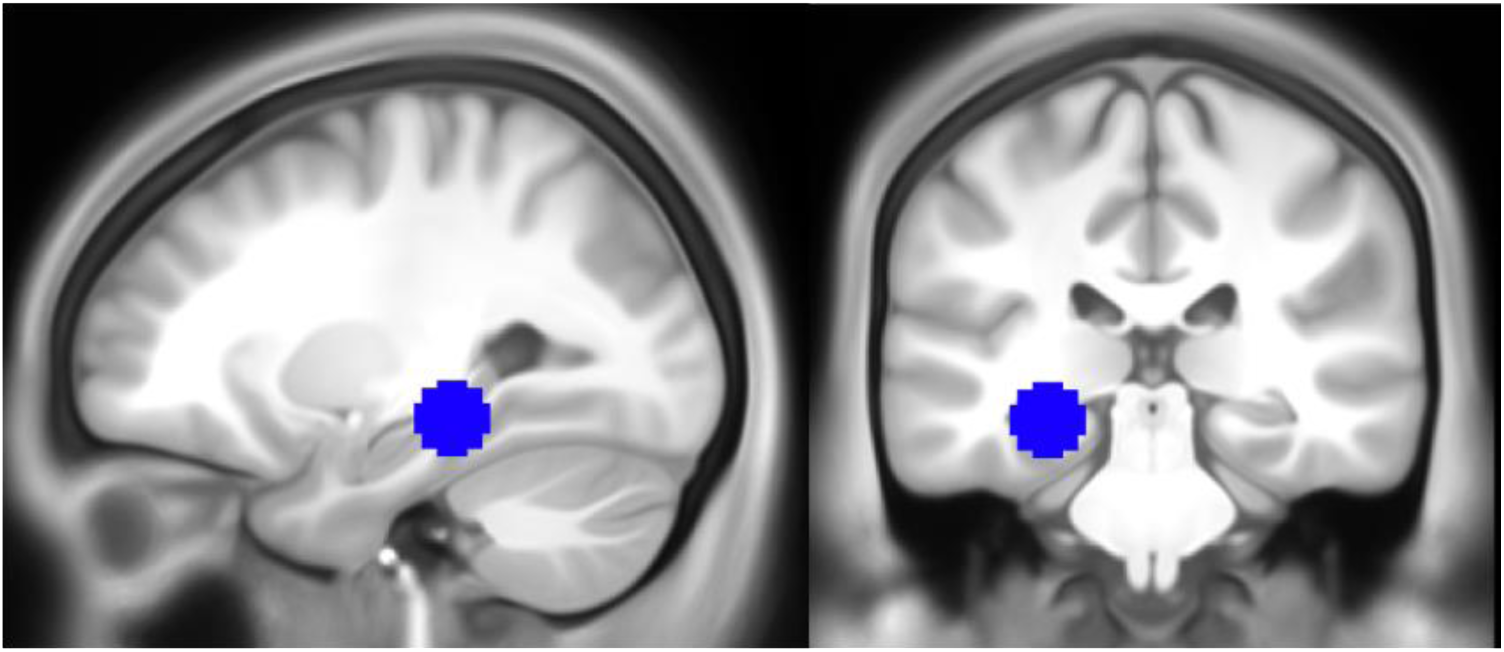
The hippocampal ROI used in the MVPA and RSA, based on MNI coordinates extracted from Bergouignan et al. (2014; X= -27, Y = -31, Z = -11).

#### Representational Similarity Analysis: Encoding-Retrieval Pattern Similarity

We employed representational similarity analysis (RSA) to investigate whether the condition (synchronous or asynchronous) associated with each video was related to the degree of similarity between spatial patterns of BOLD activity during memory encoding and retrieval (i.e. encoding-retrieval similarity) at the wholebrain and ROI level. We carried out additional wholebrain and ROI analyses to test whether type of a stronger body ownership illusion and memory vividness interacted to influence encoding-retrieval similarity.

First, we created subject-specific contrast matrices designed to assess whether correlations between patterns of activity associated with memory encoding and retrieval were higher in the synchronous compared to asynchronous condition (see Figure 2A). Normalized beta estimates from each video across the three repetitions in each session (i.e., encoding, retrieval) were averaged together, resulting in 24 beta estimates per session (i.e. 12 per condition, 1 per video), and used as input to the analysis. Videos associated with the synchronous condition were weighted positively while videos associated with the asynchronous condition were weighted negatively, such that the diagonal of the contrast matrix sums to zero. This analysis sought to identify brain regions where a stronger body ownership illusion (synchronous) led to greater memory reinstatement, relative to asynchronous condition with a supressed illusion. For example, if the video for Moderna Museet was encoded with synchronous visuotactile stimulation, then this video received a weighting of positive one. In contrast, if the video for Central Station was encoded with asynchronous visuotactile stimulation, then it received a weighting of negative one.

**Figure 2.**
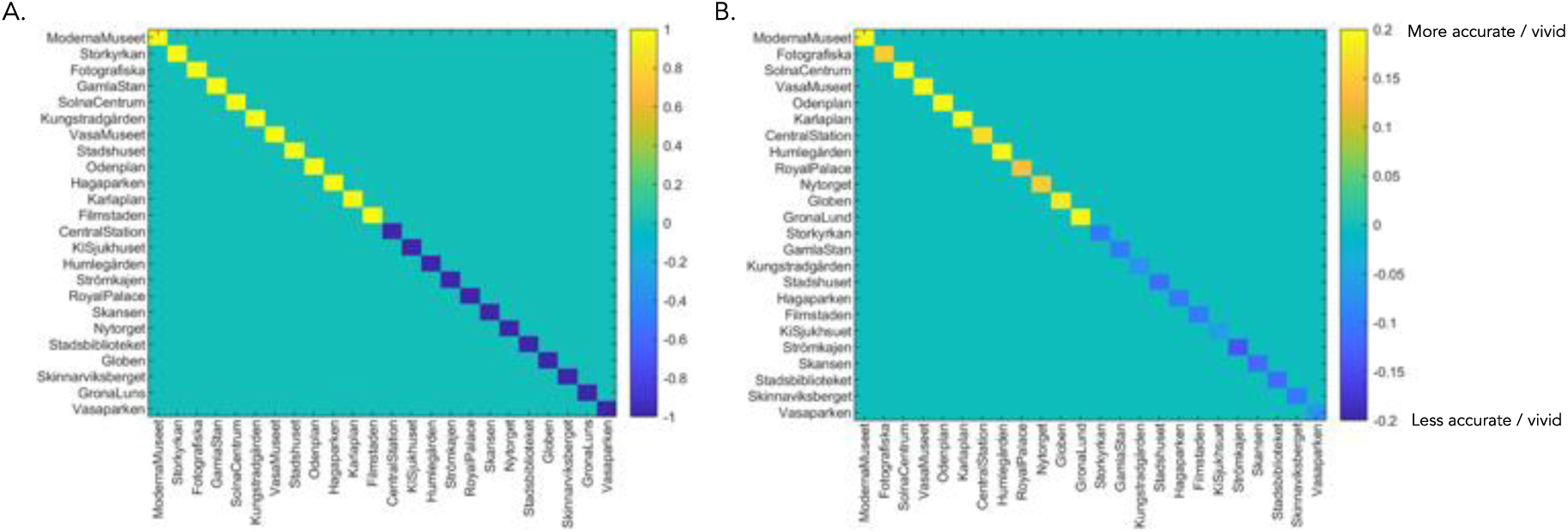
*RSA Contrast Matrices*. (A) We created individual subject contrast matrices comparing pattern similarity between encoding and retrieval of the same video where similarity is higher in the synchronous condition (on-diagonal). Videos encoded with the body ownership illusion (synchronous condition) received positive weights (i.e. yellow on-diagonal values), while videos encoded with reduced illusion (asynchronous) received negative weights (i.e. blue on-diagonal values) (B) Same-video correlations (on-diagonal) were weighted according to condition (i.e. synchronous = positive/yellow on-diagonal values, asynchronous = negative/blue on-diagonal values) and factor of interest (e.g. memory accuracy). This RSA detects regions where reinstatement is greater for videos encoded with the body ownership illusion and increasing memory richness. An example contrast matrix from a single participant is shown here.

Second, we performed an additional type of analysis where we weighted correlations positively or negatively by convolving condition (i.e. synchronous = positive, asynchronous = negative) and memory vividness, as measured by the average vividness reported post-scanning on session one and session two (see Figure 2B; (Oedekoven et al., 2017). For example, if the video for Moderna Museet were encoded with synchronous visuotactile stimulation and received a vividness rating of four, then this video received a contrast value of five (i.e. vividness rating plus the value for the synchronous condition), which was divided by the sum of the contrast values for all videos such that the contrast values for all videos summed to one. For a video encoded with asynchronous visuotactile stimulation, a value of one would be subtracted rather than added to a participant’s vividness rating before it was divided by the sum of the contrast values for all videos. Contrast values corresponding to each video were assigned to the diagonal of the participant’s contrast matrix, and all off-diagonal values were set to sum to negative one, such that the final contrast matrix totalled zero. This analysis identified areas where the degree of encoding-retrieval pattern similarity scaled with the body ownership illusion (synchronous vs. asynchronous) and the vividness of retrieval.

For the wholebrain analyses, a spherical searchlight of 100 voxels was centred at each voxel in the acquired volumes. The summed difference in Fisher-transformed Pearson correlations between encoding and retrieval of the same video was assigned to the centre voxel of the searchlight. The individual searchlight maps were analyzed at the group level by submitting them to a one-sample t-test corrected for multiple comparisons using TFCE (Smith & Nichols, 2009) with 10,000 Monte Carlo simulations to identify regions where correlations were statistically greater than zero (e.g. Oedekoven et al., 2017). For the ROI analyses, we tested whether the body ownership illusion (synchronous versus asynchronous), and its interaction with memory vividness during retrieval, influenced encoding-retrieval pattern similarity in 1) the hippocampus, and 2) brain regions that were sensitive to encoding memories with a stronger body ownership illusion (synchronous versus asynchronous) as measures of memory reinstatement. We used the same hippocampal ROI as in the multivariate pattern analysis of the encoding and retrieval sessions. We created the ROI for the regions able to decode synchronicity of visuotactile stimulation during encoding based on the group-level results of the MVPA, which was converted into binary format using the MarsBar toolbox for SPM (Brett, 2002) and resampled to 2mm cubic voxels. For both ROIs, encoding-retrieval pattern similarity was estimated using Pearson correlations and compared to the relevant contrast matrix for each participant. Subject-level correlations were then Fisher-transformed to approximate a normal distribution more closely for statistical analysis, and submitted to one-sided, one-sample t-tests against a null hypothesis of zero. The t-tests were one-sided because we predicted that the average correlations would be greater, but not less, than zero.

#### Univariate Analysis of the Encoding and Retrieval Sessions

To confirm that our experimental design activated expected memory encoding and retrieval regions, regardless of type of visuotactile congruence, we conducted univariate analyses for each fMRI session separately. These analyses were a sanity check to ensure that our experimental design activated brain regions commonly implicated in each phase of memory processing. We used the same trial-wise beta-estimates from the encoding and retrieval sessions as in the multivariate analyses, except that they were spatially smoothed with a 8mm full-width at half maximum Gaussian kernel, as opposed to a 3mm full-width at half maximum Gaussian kernel, consistent with standards for univariate analyses (Mikl et al., 2008). Further, in line with recent studies (for review see D’Argembeau et al., 2022) and temporal compression effects in autobiographical memory (e.g. Jeunehomme et al., 2020), the analysis of the retrieval session was constricted to the first ten seconds of each trial.

Memories involving the retrieval of specific performed actions increase event segmentation and decrease temporal compression (Jeunehomme et al., 2020). Since the memories in the present study involved the participant passively watching the scene without performing a physical action and consisted of only one main narrative event (i.e. limited event segmentation), we reasoned that the rate of temporal compression would be greater than those reported by previous studies using autobiographical memories. Thus, we chose a compression factor of two (i.e. twenty seconds of encoding shortened to ten seconds of retrieval). For the encoding session, beta weights reflecting the effects of forming memories for each condition (i.e. synchronous, asynchronous) were contrasted against baseline estimates for each participant. For the retrieval session, beta weights that reflected the first ten seconds of memory retrieval were contrasted against baseline estimates for each participant. Group-level effects were estimated with the Threshold Free Cluster Enhancement toolbox for SPM12 (Gaser, 2019) using 10,000 permutations and corrected for multiple comparisons with false discovery rates (*q* < .05).

## Behavioural Results

### Questionnaire Responses

We performed a paired-samples t-test comparing average illusion minus average control ratings for the synchronous and asynchronous condition separately to assess the subjective strength of the illusion induction. Critically, difference ratings were significantly higher in the synchronous (*M* = 3.13, *SD* = 1.52) compared to asynchronous condition (*M* = 2.31, *SD* = 1.87), *t*(29) = 2.08, *p* = .047, Cohen’s d = .38).^1^ Consistent with this finding, a direct comparison of average illusion statement ratings between the synchronous (*M* =1.19, *SD* = .24) and asynchronous visuotactile condition (*M* = .23, *SD* = 1.78) was also significant, *t*(29) = 2.55, *p* = .02, Cohen’s d = .47 (see Figure 3A). These findings confirm that the participants felt a stronger illusion of embodying the mannequin following synchronous compared to asynchronous visuotactile stimulation as expected (Guterstam, Björnsdotter, Gentile, & Ehrsson, 2015; Iriye & Ehrsson, 2022; Petkova et al 2008; O’Kane & Ehrsson, 2021).^1^

**Figure 3.**
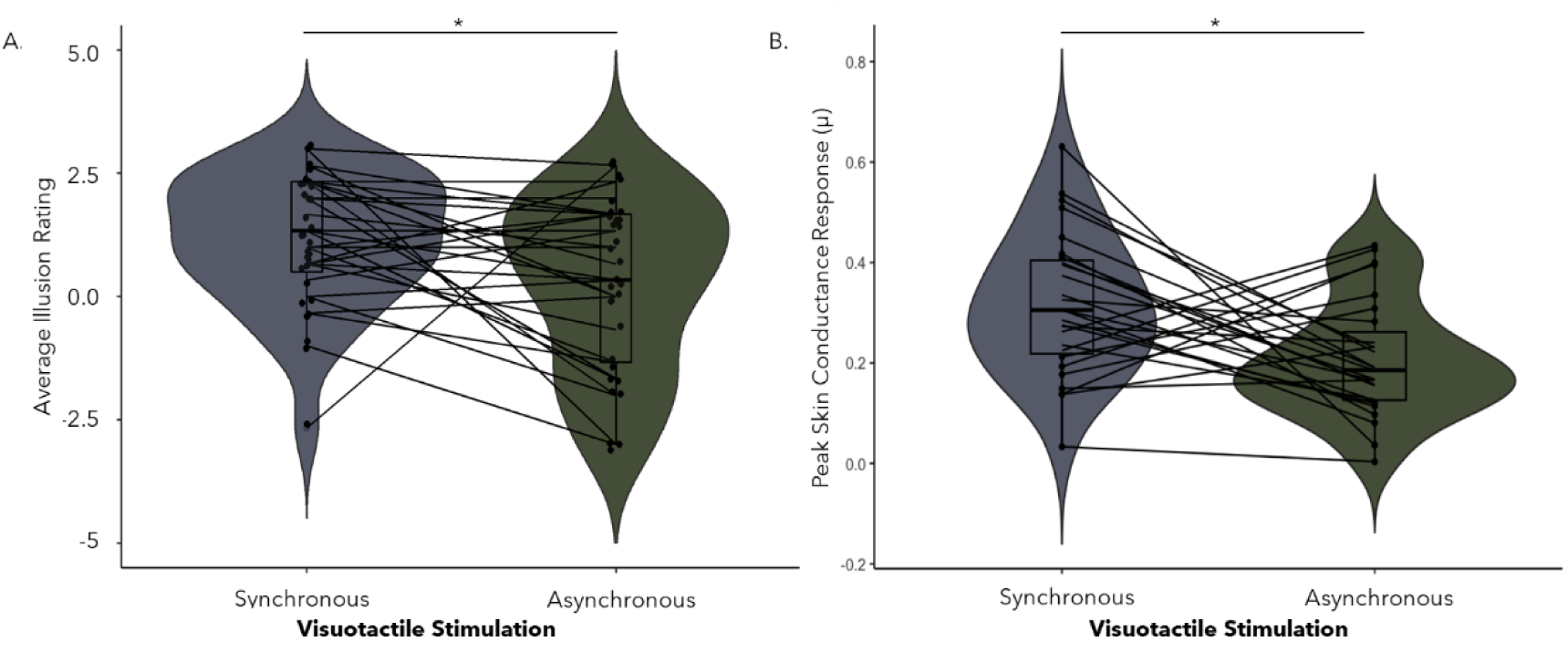
(A) The average illusion statement scores was significantly higher following synchronous compared to asynchronous visuotactile stimulation, *p* = .02 (A). (B) The peak magnitudes of skin conductance responses were higher for knife threats experienced during the synchronous compared to asynchronous condition, *p* = .01.

### Skin Conductance Responses

A paired-samples t-test comparing SCR magnitudes between conditions found a significant effect of visuotactile congruence (*t*(25) = 2.69, *p* = .01, Cohen’s *d* = .53).^2^ SCR magnitude was higher in the synchronous (*M* = .32, *SD* = .15) compared to asynchronous visuotactile stimulation condition (*M* = .20, *SD* = .12, see Figure 3B). This finding provides physiological evidence that illusory ownership over the mannequin’s body was higher in the synchronous compared to asynchronous condition (Petkova et al., 2008; Guterstam et al., 2015).

### Eye Tracking Data

Average fixation coordinates were plotted for both experimental conditions for each participant (see Figure 4A&B). A 2 (visuotactile stimulation: synchronous, asynchronous) x 2 (fixation coordinate: X,Y) did not reveal a significant main effect of fixation location according to condition, nor an interaction between visuotactile congruence and fixation coordinate (*p*’s > 0.22; see Figure 6).^3^ Thus, the average gaze location did not differ according to condition, implying that eye-movements and overt attention was reasonably matched between condition. There was a significant main effect of coordinate, *F*(1,27) = 10.00, *p* = .004, η_p_^2^ = .27, but this is irrelevant as there was no interaction with congruence of visuotactile stimulation.

**Figure 4.**
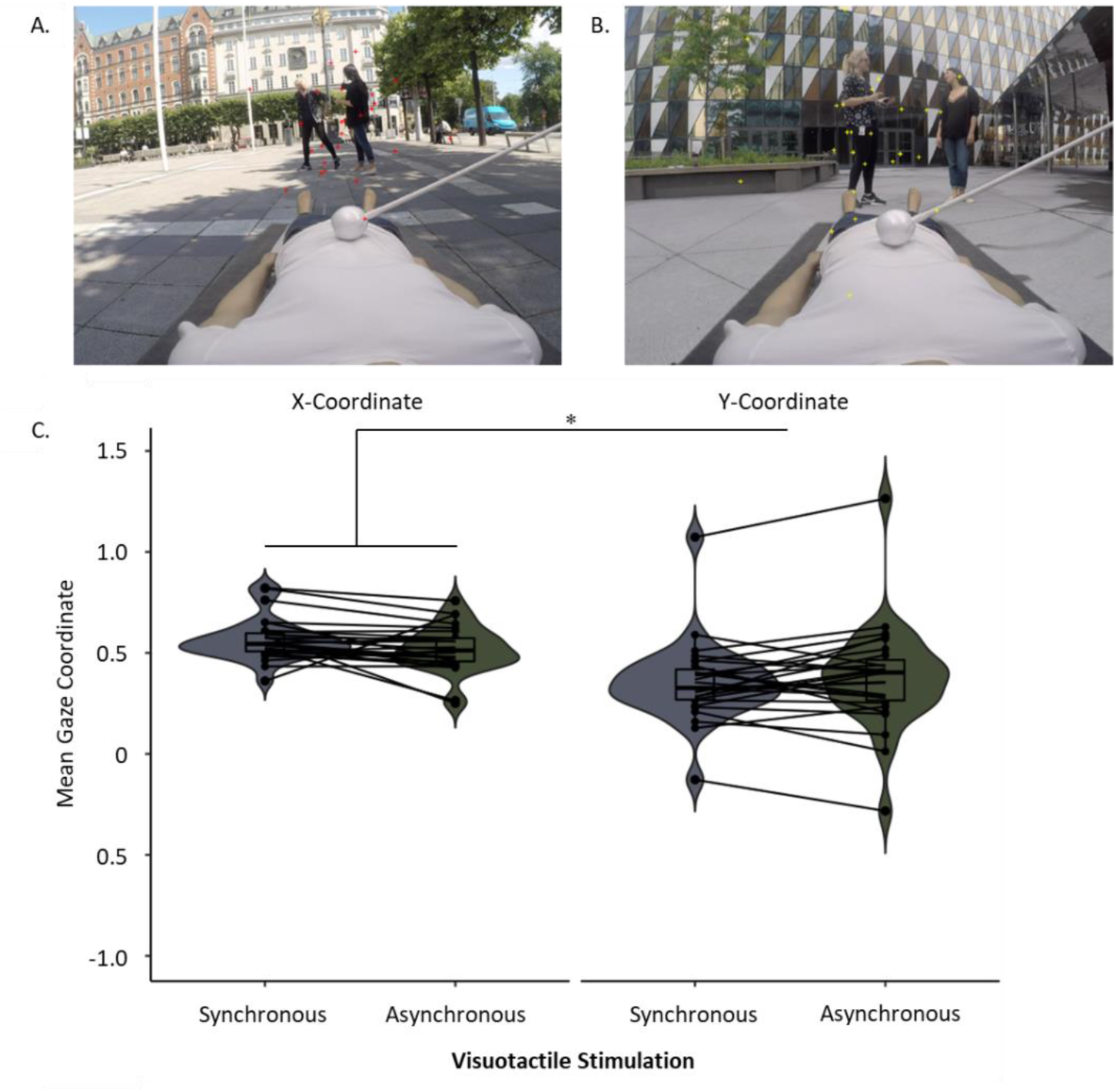
Average fixation coordinates did not differ between the synchronous (A) and asynchronous (B) conditions. Each cross represents the average fixation across the video for one participant. There was no main effect of type of visuotactile stimulation on mean gaze (*p*=0.22), or interaction between type of visuotactile stimulation and co-ordinate (*p* = .19; C).

**Figure 5.**
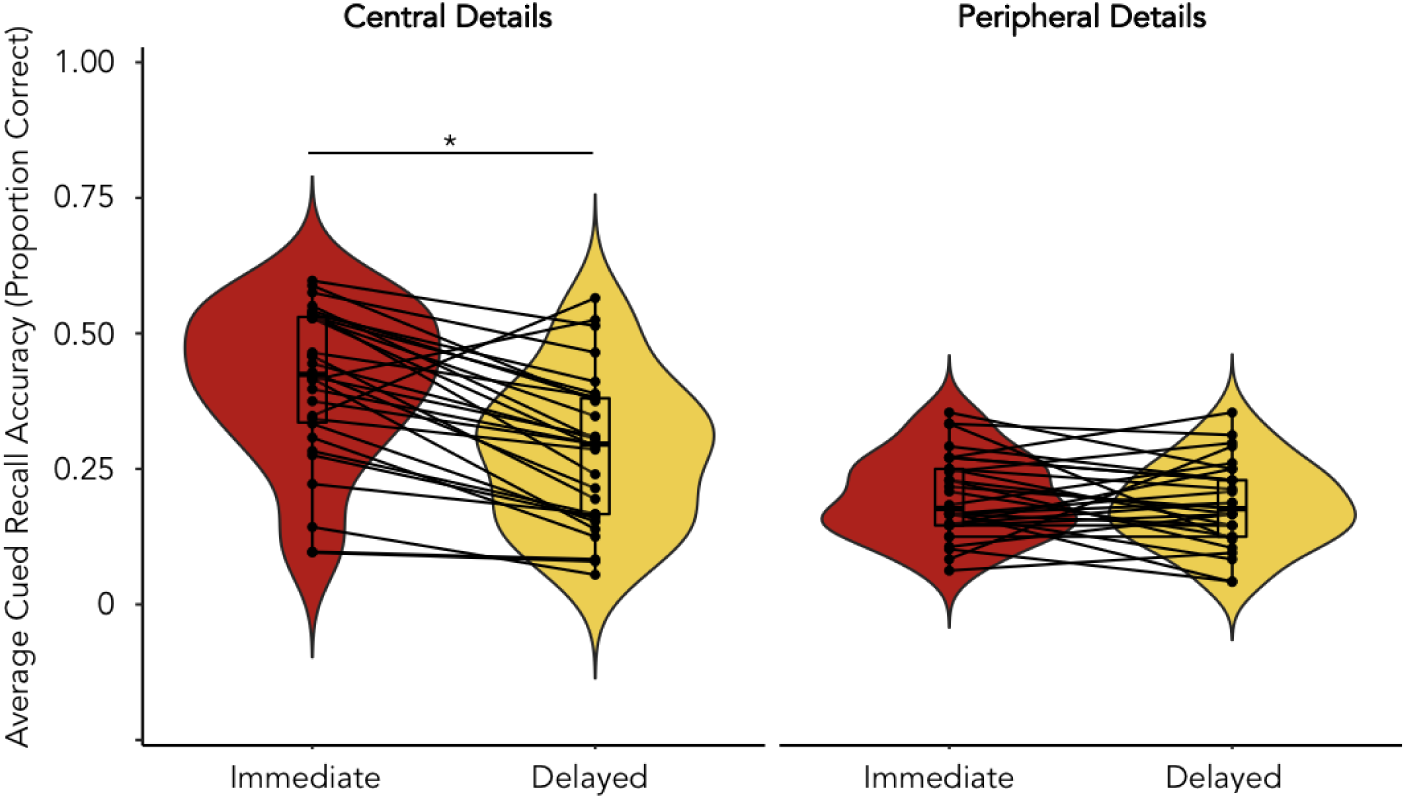
Average cued recall accuracy was higher at immediate testing compared to delayed testing for central event details (*p* < .001), but not peripheral details.

**Figure 6.**
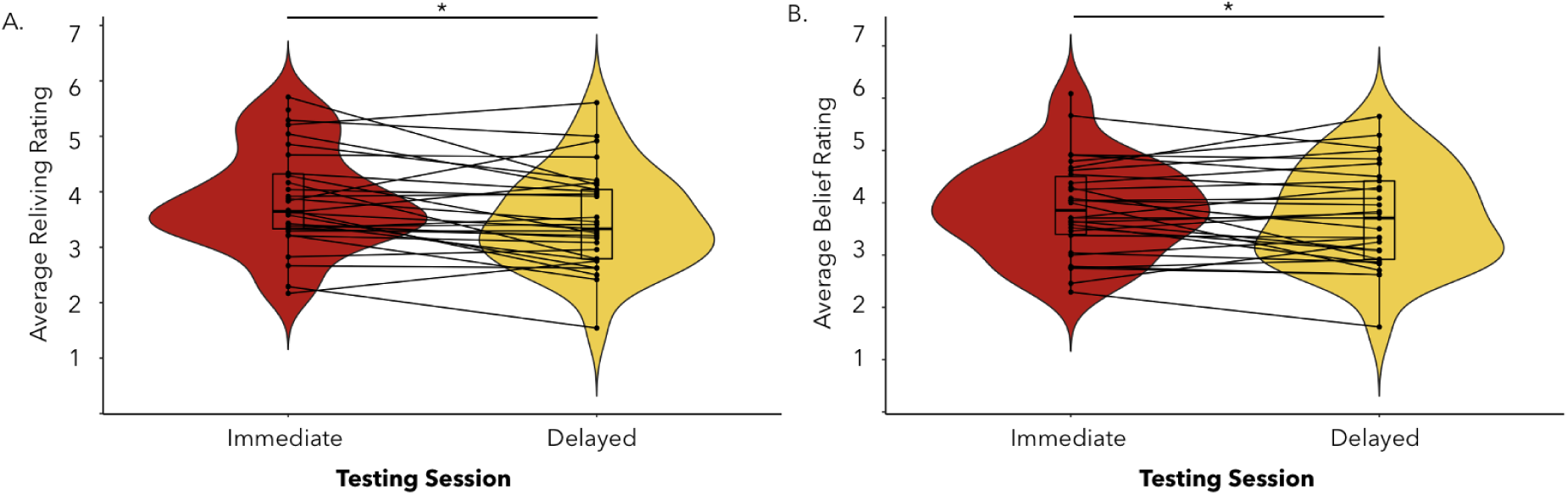
Average reliving (A) and belief in memory accuracy (B) ratings were higher at immediate compared to delayed testing, *p*’s < .02.

### Presence

Ratings from the three presence statements were averaged together separately for the synchronous (*M* = 3.38, *SD* = 1.14) and asynchronous (*M* = 3.64, *SD* = 1.29) visuotactile congruence conditions. A paired sample t-test did not reveal any significant differences between conditions, *F*(1,29) = 1.32, *p* = .26, Cohen’s *d* = .04, demonstrating that both types of visuotactile stimulation conditions induced similar levels of presence within the immersive videos.^4^

### Cued Recall Accuracy

A 2 (testing session: immediate, delayed) x 2 (detail type: central, peripheral) x 2 (visuotactile congruence: synchronous, asynchronous) repeated measures ANOVA revealed significant main effects of testing session, *F*(1,28) = 27.10, *p* < .001, η_p_^2^ = .49, and detail type, *F*(1,28) = 49.84, *p* < .001, η_p_^2^ = .64, but not visuotactile congruence *F*(1,28) = 0.01, *p* = 0.92, . η_p_^2^ = 0.0004. Main effects were qualified by a significant interaction between testing session and detail type, *F*(1,28) = .78, *p* < .001, η_p_^2^ = .39. Post-hoc tests with Bonferroni corrections indicated that memory accuracy for central details decreased between testing points (immediate: *M* = .41, *SD* = .14; delayed: *M* = .28, *SD* = .14; see Figure 5), *p* < .001. In contrast, memory accuracy for peripheral details did not change between testing points (immediate: *M* = .20, *SD* = .08; delayed: *M* = .18, *SD* = .18, see Figure 5), *p* = 1.00.^5^

### Subjective Ratings

We conducted four separate 2 (testing session: immediate, delayed) x 2 (visuotactile congruence: synchronous, asynchronous) repeated measures ANOVAs on participants’ ratings of vividness, reliving, perceived memory accuracy, and emotional intensity. For vividness ratings, there was a significant interaction between session and type of visuotactile congruence, *F*(1,27) = 4.51, *p* = .04, η_p_^2^= .14. However, follow-up post-hoc tests with Bonferroni corrections did not reveal any significant effects, all *p*’s > .63.^6^ There was a main effect of testing session on reliving ratings, *F*(1,27) =6.00, *p* = .02, η_p_^2^= .18, which indicated higher reliving at immediate (*M* = 3.86, *SD* = .91) compared to delayed testing (*M* = 3.51, *SD* = .91; see Figure 6A).^6^ There was also a main effect of testing session on perceived accuracy ratings, *F*(1,27) = 15.03, *p* < .001, η_p_^2^= .36, whereby participants believed in the accuracy of their memories to a higher degree during immediate (*M* = 3.85, *SD* = .94) relative to delayed testing (*M* = 3.35, *SD* = 1.06; see Figure 6B).^7^ For emotional intensity ratings, there were no significant main effects or interactions.^8^

## fMRI Results

### Multivariate Analysis of Memory Encoding

A wholebrain multivariate pattern analysis of the period of memory encoding during session one revealed a distributed set of regions where patterns of neural activity could distinguish between the synchronous and asynchronous conditions, thereby decoding the full-body illusion, i.e., whether body ownership of the mannequin was stronger or weaker (Table 3). The identified regions included the left ventral premotor cortex, right intraparietal sulcus, bilateral lateral occipital cortex (Figure 7), and putamen, and lateral cerebellum, which have been previously linked to feelings of body ownership in full-body illusion paradigms (Guterstam et al., 2015; Preston & Ehrsson 2016; Petkova et al., 2011) and rubber hand illusion studies (Ehrsson et al., 2004; Gentile et al., 2013; Limanowski et al., 2016) . These findings thus confirms that the participants were experiencing a stronger illusion in the synchronous condition than in the asynchronous condition in line with the questionnaire and threat-evoked results reported above.

**Figure 7.**
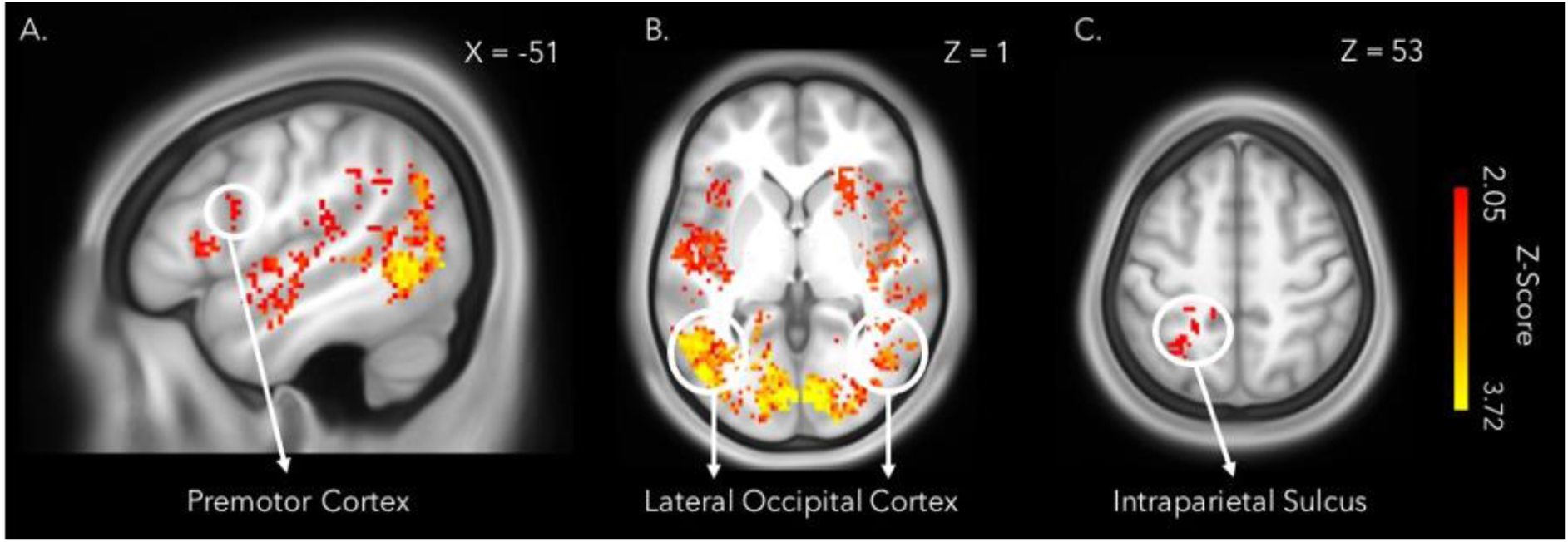
The wholebrain searchlight analysis decoding illusion condition (synchronous or asynchronous) during encoding of the immersive videos identified several regions previously implicated in the sense of body ownership, including the left premotor cortex (A), bilateral lateral occipital cortex (B), and the left intraparietal sulcus (C). The statistical map was thresholded at *FDR* < .05 and overlaid onto the average T1w images of 440 subjects obtained from the WU-Minn HCP dataset (van Essen et al., 2013).

**Table 3.**

| Table 3. MVPA Encoding Session |  |  |  | MNI Coordinates |  |  |
| --- | --- | --- | --- | --- | --- | --- |
| Anatomical Region | BA | Extent | q-value | x | y | z |
| Orbitofrontal Cortex | 11 | 328 | 2,447 | 22 | 26 | -16 |
| Inferior Frontal Gyrus, Pars Opercularis | 44 | 478 | 2,280 | -54 | 6 | 16 |
|  | 44 | 1 | 2,395 | 52 | 18 | 0 |
| Inferior Frontal Gyrus, Pars Orbitalis | 47 | 2 | 2,409 | 38 | 24 | -12 |
| Inferior Frontal Gyrus, Pars Triangularis | 46 | 478 | 2,848 | -38 | 20 | 4 |
|  | 46 | 478 | 2,512 | -44 | 30 | 22 |
|  | 45 | 32 | 2,512 | 46 | 22 | 6 |
|  | 45 | 1 | 2,395 | 38 | 30 | 6 |
| Rolandic Operculum | 6 | 2 | 2,152 | 46 | -2 | 20 |
| Insula | 13 | 826 | 2,863 | -40 | -4 | 2 |
|  | 13 | 5 | 2,160 | 34 | -18 | 24 |
|  | 13 | 3 | 2,644 | 40 | -10 | 8 |
|  | 13 | 2 | 2,409 | 36 | -20 | 8 |
|  | 13 | 1 | 2,697 | 34 | -8 | 10 |
|  | 13 | 1 | 2,395 | 46 | 14 | 0 |
|  | 13 | 1 | 2,223 | -34 | 16 | -10 |
|  | 13 | 1 | 2,223 | -38 | -4 | -8 |
|  |  | 1 | 2,152 | -36 | 12 | 6 |
| Heschl's Gyrus | 41 | 826 | 2,834 | -36 | -26 | 8 |
|  | 41 | 4 | 2,848 | 34 | -32 | 14 |
|  | 41 | 1 | 2,165 | -62 | -28 | 12 |
| Temporal Pole | 38 | 6 | 2,301 | -40 | 8 | -16 |
|  | 38 | 1 | 2,160 | 46 | 8 | -20 |
| Hippocampus | 36 | 1 | 2,644 | 32 | -16 | -20 |
| Parahippocampal Gyrus |  | 1 | 2,395 | 30 | -34 | -12 |
| Superior Temporal Gyrus | 22 | 826 | 2,794 | -60 | -14 | 10 |
|  | 22 | 31 | 2,423 | 60 | -16 | 0 |
|  | 22 | 9 | 2,782 | 62 | -22 | -8 |
|  | 22 | 3 | 2,707 | 62 | -14 | -6 |
|  | 22 | 3 | 2,414 | 44 | -2 | -16 |
|  | 22 | 1 | 2,223 | -44 | -26 | 0 |
|  | 22 | 1 | 2,170 | 44 | -34 | 8 |
|  | 22 | 1 | 2,152 | 44 | -30 | -2 |
|  | 22 | 1 | 2,044 | -50 | -40 | 22 |
| Middle Temporal Gyrus | 21 | 42 | 2,350 | -64 | -38 | 10 |
|  | 21 | 3 | 2,160 | -48 | -32 | -4 |
|  | 21 | 1 | 2,644 | 60 | -40 | -4 |
|  | 21 | 1 | 2,223 | -38 | -30 | 0 |
|  | 21 | 1 | 2,152 | 62 | -32 | 6 |
| Inferior Temporal Gyrus | 38 | 1 | 2,549 | -46 | -4 | -28 |
| Premotor Cortex | 6 | 2 | 2,152 | 38 | -2 | 26 |
|  | 6 | 1 | 2,152 | 40 | -8 | 28 |
| Precentral Gyrus | 4 | 1 | 2,432 | 42 | -4 | 16 |
| Postcentral Gyrus | 1 | 1 | 2,152 | 36 | -12 | 28 |
|  | 1 | 1 | 2,152 | 50 | -16 | 32 |
| Dorsal Posterior Cingulate Cortex | 31 | 14 | 2,162 | -12 | -38 | 50 |
|  | 31 | 2 | 2,062 | -6 | -38 | 50 |
| Precuneus | 7 | 2 | 2,167 | 6 | -70 | 46 |
| Table 3. continued |  |  |  |  |  |  |
| Superior Parietal Gyrus | 7 | 101 | 2.391 | -28 | -52 | 52 |
|  | 7 | 4 | 2.062 | -18 | -46 | 48 |
|  | 7 | 3 | 2.054 | -16 | -62 | 58 |
|  | 7 | 1 | 2.115 | -24 | -56 | 58 |
|  | 7 | 1 | 2.068 | -28 | -62 | 48 |
|  | 7 | 1 | 2.068 | -18 | -62 | 54 |
|  | 7 | 1 | 2.046 | -30 | -58 | 30 |
|  | 7 | 1 | 2.044 | -22 | -44 | 46 |
|  | 7 | 1 | 2.044 | -16 | -66 | 54 |
| Superior Parietal Sulcus | 5 | 9 | 2.162 | -24 | -36 | 52 |
|  | 5 | 2 | 2.071 | -12 | -32 | 54 |
|  | 5 | 1 | 2.068 | -16 | -36 | 58 |
| Angular Gyrus | 39 | 58 | 2.197 | -46 | -50 | 32 |
|  | 39 | 10 | 2.135 | -34 | -52 | 24 |
|  | 39 | 6 | 2.086 | -40 | -58 | 32 |
|  | 39 | 4 | 2.046 | -36 | -58 | 32 |
|  | 39 | 3 | 2.104 | -32 | -60 | 38 |
|  | 39 | 2 | 2.058 | -54 | -48 | 34 |
|  | 39 | 1 | 2.409 | -50 | -60 | 38 |
|  | 39 | 1 | 2.160 | 48 | -36 | 24 |
|  | 39 | 1 | 2.073 | -54 | -42 | 32 |
|  | 39 | 1 | 2.068 | -38 | -56 | 48 |
|  | 39 | 1 | 2.056 | -28 | -62 | 38 |
|  | 39 | 1 | 2.048 | -50 | -48 | 36 |
| Supramarginal Gyrus | 40 | 49 | 2.257 | -48 | -30 | 12 |
|  | 40 | 27 | 2.214 | -36 | -32 | 16 |
|  | 40 | 18 | 2.512 | 62 | -16 | 22 |
|  | 40 | 3 | 2.294 | -48 | -24 | 10 |
|  | 40 | 3 | 2.175 | 62 | -18 | 10 |
|  | 40 | 3 | 2.062 | -56 | -44 | 30 |
|  | 40 | 1 | 2.782 | 62 | -22 | 20 |
|  | 40 | 1 | 2.395 | 50 | -20 | 28 |
|  | 40 | 1 | 2.152 | 60 | -16 | 30 |
|  | 40 | 1 | 2.066 | -52 | -38 | 26 |
|  | 40 | 1 | 2.062 | -56 | -40 | 34 |
|  | 40 | 1 | 2.062 | -58 | -38 | 38 |
| Middle Occipital Gyrus | 19 | 4 | 2.468 | -42 | -76 | 26 |
|  | 19 | 2 | 2.160 | -34 | -64 | 10 |
|  | 19 | 1 | 2.160 | -34 | -68 | 14 |
| Fusiform Gyrus | 37 | 4 | 2.162 | -32 | -54 | 2 |
|  | 37 | 1 | 2.782 | -36 | -48 | -16 |
|  | 37 | 1 | 2.167 | -44 | -46 | -10 |
|  | 37 | 1 | 2.160 | 32 | -66 | -16 |
| Lingual Gyrus | 18 | 8526 | 3.719 | -20 | -80 | -12 |
| cluster extends to the left parahippocampal gyrus |  |  |  |  |  |  |
| cluster extends to the right hippocampus | 18 | 8526 | 3.719 | 14 | -86 | -8 |
|  | 18 | 3 | 2.106 | 18 | -52 | 2 |
|  | 18 | 2 | 2.409 | -20 | -52 | 4 |
|  | 18 | 2 | 2.104 | 20 | -56 | 0 |
|  | 17 | 1 | 3.036 | 26 | -74 | 4 |
|  | 17 | 1 | 2.929 | 28 | -72 | 2 |
|  |  | 1 | 2.152 | 24 | -88 | -8 |
|  | 19 | 1 | 2.084 | 18 | -58 | -4 |
|  | 19 | 1 | 2.395 | 26 | -44 | -8 |
| Table 3. continued |  |  |  |  |  |  |
| Calcarine Sulcus | 19 | 8526 | 3.719 | -40 | -80 | -10 |
|  | 19 | 8 | 2.794 | -28 | -64 | 14 |
|  | 17 | 5 | 2.223 | 30 | -68 | 10 |
|  | 17 | 1 | 2.644 | -16 | -80 | 12 |
|  | 17 | 1 | 2.395 | 18 | -74 | 14 |
|  | 17 | 1 | 2.152 | -20 | -64 | 18 |
| Caudate | - | 2 | 2.457 | 18 | 26 | 14 |
| Putamen | - | 328 | 2.848 | 26 | 18 | 4 |
|  | - | 328 | 2.437 | 30 | 2 | -8 |
|  | - | 16 | 2.748 | 32 | 2 | 4 |
|  | - | 1 | 2.644 | 28 | -8 | 14 |
|  | - | 1 | 2.395 | 26 | -6 | 10 |
|  | - | 1 | 2.160 | 24 | 12 | 6 |
|  |  | 1 | 2.152 | 22 | 10 | 4 |
| Thalamus |  | 1 | 2.418 | 16 | -34 | 8 |
| Cerebellum Vermis (IV/V) |  | 1 | 2.395 | -2 | -58 | -4 |
| Cerebellum Vermis (VI) | - | 1 | 2.782 | -2 | -64 | -18 |
| Cerebellum Crus 1 Lobule | - | 6 | 2.506 | -22 | -66 | -30 |
|  |  | 3 | 2.669 | -38 | -72 | -24 |
|  |  | 3 | 2.395 | 34 | -52 | -36 |
|  | - | 2 | 2.160 | 34 | -58 | -34 |
| Cerebellum Lobule VI | - | 135 | 2.506 | 14 | -70 | -16 |
|  | - | 14 | 2.437 | 30 | -46 | -30 |
|  | - | 2 | 2.782 | 30 | -52 | -22 |
|  |  | 1 | 2.418 | -22 | -66 | -14 |
|  |  | 1 | 2.395 | 32 | -50 | -26 |
|  | - | 1 | 2.175 | 14 | -74 | -24 |

More importantly for the main questions of the present study, the whole-brain analysis further revealed a set of medial temporal and ventral posterior parietal regions known to be essential to successful memory encoding, including the right hippocampus, bilateral parahippocampal gyri (extending into the fusiform gyri), and bilateral angular gyri (Figure 8). Lateral frontal and temporal regions were also sensitive to the full-body illusion during encoding. While patterns of activity in the left hippocampus did not survive corrections for multiple comparisons at the whole-brain level, the ROI analysis revealed that the same hippocampal region previously identified by Bergouignan and colleagues (2014) could significantly predict whether a memory was encoded with illusory ownership (synchronous) or not (asynchronous), *t*(23) = 2.58, *p* = .02, Cohen’s *d* = (*M* = .54, *SD* = .07).

**Figure 8.**
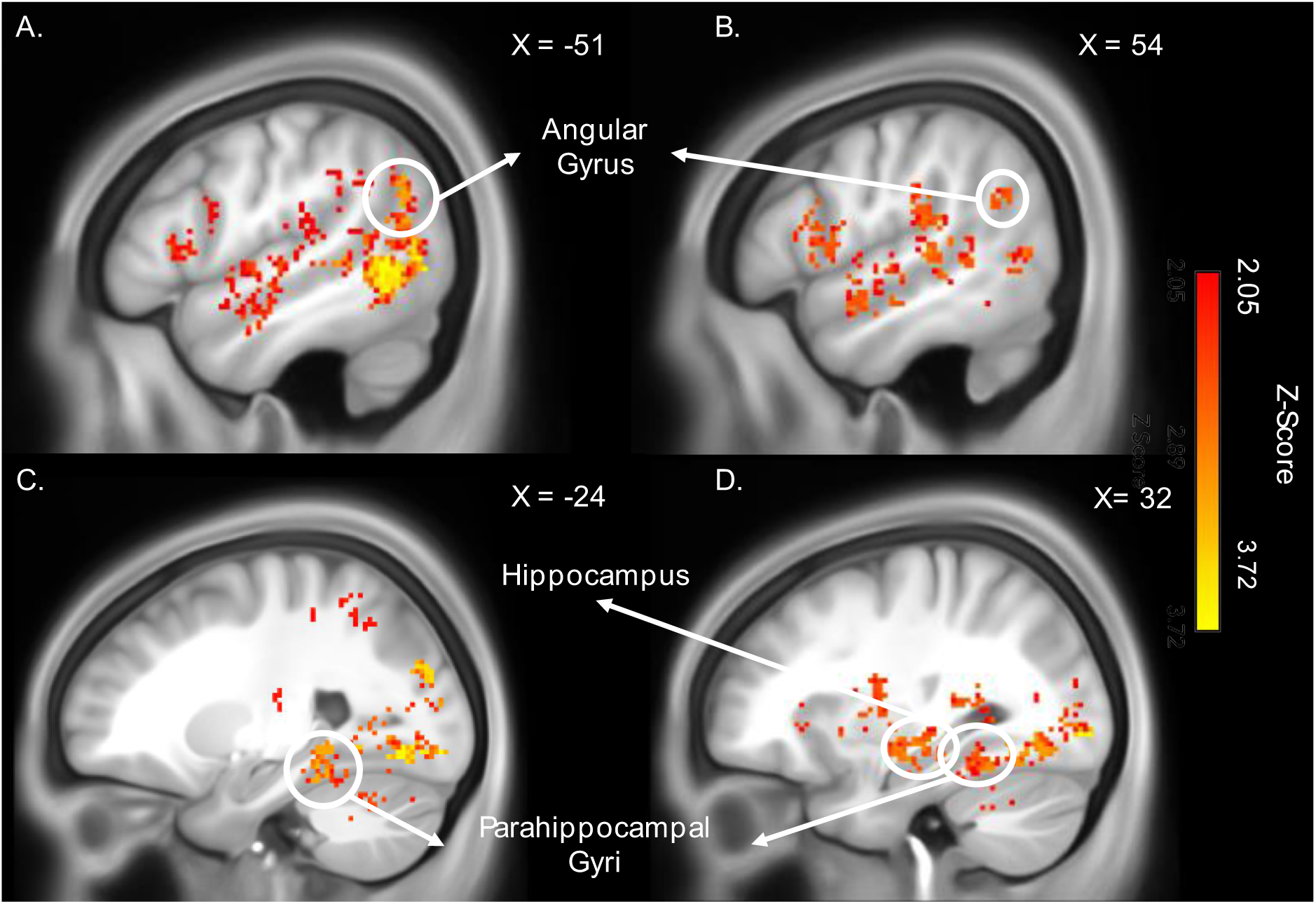
Patterns of activity in posterior parietal and medial temporal lobe regions known to be crucial to successful memory formation predicted body illusion condition (synchronous or asynchronous) during the encoding of the immersive videos in a wholebrain searchlight analysis. The regions identified included bilateral angular gyri (A&B), the right hippocampus (C), and bilateral parahippocampal gyri (C&D). The statistical map was thresholded at *FDR* < .05 and overlaid onto the average T1w images of 440 subjects obtained from the WU-Minn HCP dataset (van Essen et al., 2013).

### Memory Retrieval

A wholebrain multivariate pattern analysis of the period of memory retrieval during session one did not reveal any regions that could significantly decode between memories that had been encoded in the synchronous and asynchronous conditions in the preceding session. The results were not affected by whether the onset of memory retrieval was set to the presentation of the memory cue or the instruction for the participant to close their eyes and begin retrieving their memory for the specified video. The results also remained the same after regressing out videos participants had forgotten (N = 23 videos, .04% of total trials) and only including the initial 10 seconds of each trial to account for the temporal compression of retrieval. Further, classification accuracy of illusion condition during encoding (synchronous vs asynchronous) was not significantly above chance level in the hippocampal ROI when taking into account the full length of each trial, *t*(23) = 0.23, *p* = .23, Cohen’s *d* = -0.29 (*M* = .48, *SD* = .07), or restricting the analysis to the initial 10 seconds of retrieval, *t*(23) = -1.07, *p* = .30, Cohen’s *d* = -0.17 (*M* = .49, *SD* = .06) .

### RSA

The wholebrain searchlight analyses did not reveal any regions where encoding-retrieval similarity was significantly higher according to condition, or condition convolved with average vividness. Similarly, encoding-retrieval similarity was not significantly higher in the synchronous compared to asynchronous condition in the hippocampal ROI, *t*(23) = -0.05, *p* = .52, Cohen’s *d* = -0.01 (*M* = - .0005, *SD =* .0481). The ROI specifying regions whose patterns of activity could decode body illusion condition (synchronous vs asynchronous) during encoding was marginally significant, *t*(23) = 1.97, *p* = .06, Cohen’s *d* = 0.39 (*M* = 0.001, *SD* = 0.004).

However, as hypothesised, once vividness was factored into the analysis, encoding-retrieval similarity in the hippocampal ROI was higher for memories encoded with a stronger full-body illusion (synchronous) compared to a weaker illusion (asynchronous), *t*(23) = 2.11, *p* = .02, Cohen’s *d* = .43 (see Figure 9). The same was true for the ROI based on regions that decoded the synchronous versus asynchronous conditions during encoding, *t*(23) = 2.37, *p* = .01, Cohen’s *d* = 0.49 (*M* = .003, *SD* = .007). Thus, the events in the immersive 3D videos formed while experiencing the mannequin’s body in the centre of the scene as one’s own body and retrieved with high vividness led to increased memory reinstatement in the same brain regions sensitive to the full-body illusion during encoding, including the left posterior hippocampus.

**Figure 9.**
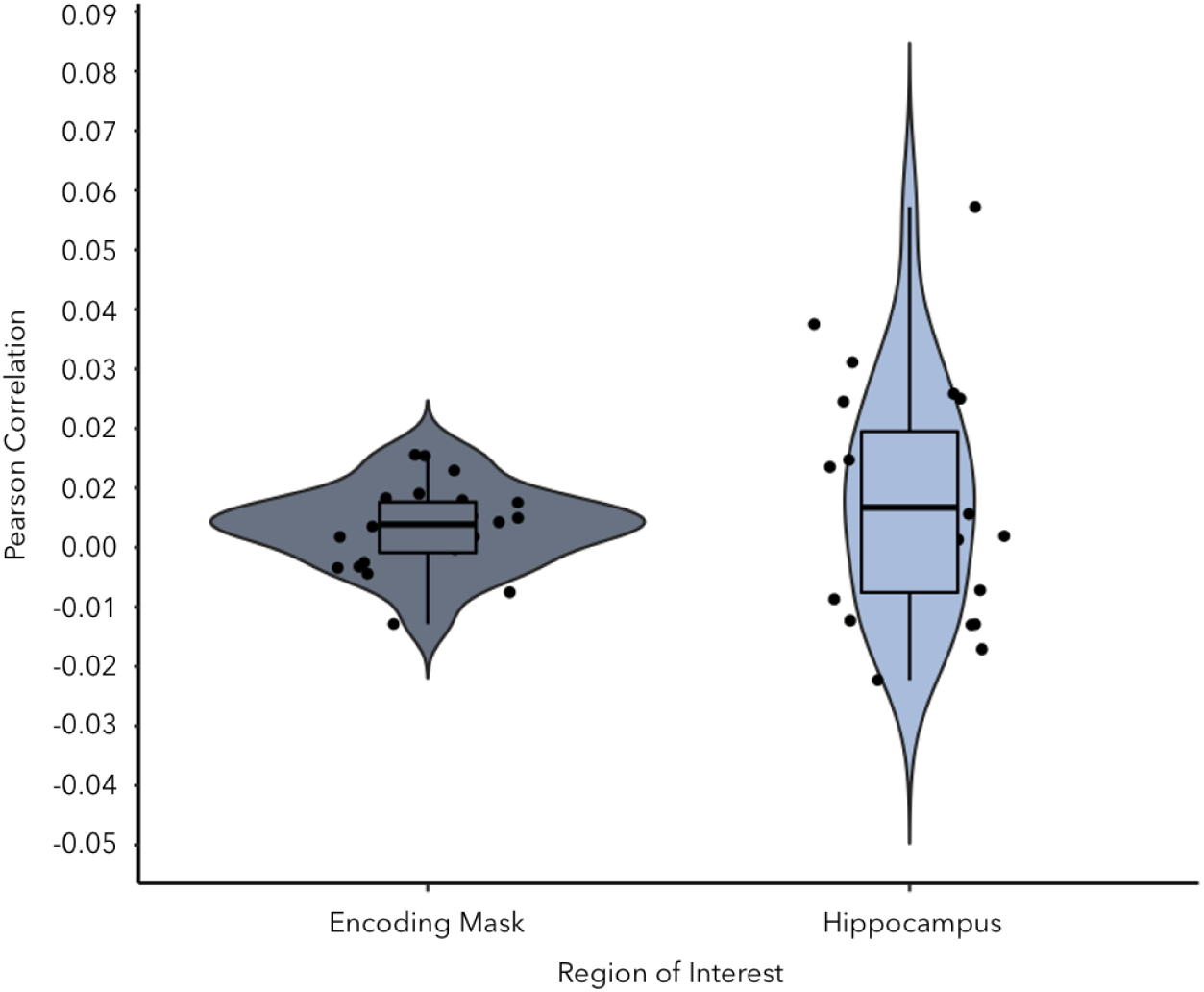
Memory reinstatement (i.e. encoding-retrieval similarity) was stronger for memories encoded with synchronous visuotactile stimulation and increasing levels of vividness in regions that decoded type of visuotactile stimulation during encoding, *p* = .01, and in the hippocampus, *p* = .02.

### Univariate Analysis of the Encoding and Retrieval Sessions

We observed activation of many regions involved in encoding the immersive naturalistic videos, regardless of synchronicity of visuotactile stimulation (see Supplemental Table 4). Large clusters of activity in the bilateral superior temporal gyri extended medially into both hippocampi, rostrally to dorsolateral prefrontal cortex, and caudally to posterior visual regions. We also identified activation in the bilateral dorsal posterior cingulate cortex and superior parietal gyrus. These findings are consistent with the previous memory encoding literature and confirms that participants were encoding the events as instructed (Gottlieb, Wong, de Chastelain & Rugg, 2012; Kim, 2015, 2019; Sonkusare et al., 2019; Spaniol et al., 2009; Rugg et al., 2015).

As expected, the retrieval of the immersive videos from memory recruited a more restricted network of brain regions which included the left dorsolateral prefrontal cortex, the left precuneus, bilateral angular gyrus, and right inferior temporal cortex (see Supplemental Tables 5). These regions are typically reported in the memory literature (Iriye & St. Jacques, 2019; Svoboda et al., 2006), and confirm that participants in our study were retrieving memories for the immersive videos as instructed.

## Discussion

The fundamental sense of our own body is at the centre of every event we experience, which affects how events are encoded and later recalled. We found that during encoding, experiencing or not experiencing a mannequin’s body as one’s own in the centre of a naturalistic and immersive multisensory scene and social event led to different patterns of activity in regions associated with memory formation, i.e, the bilateral hippocampi, parahippocampal gyri, angular gyri, the lateral prefrontal cortex, and lateral temporal cortex. Our findings demonstrate that regions critical to memory encoding form memories for events according to the degree of body ownership experienced as an event unfolds, which supports the hypothesis that bodily self-awareness is intrinsically part of episodic memories as they form. Further, reinstatement of the original memory trace was greater for memories formed with a strong compared to weak sense of illusory body ownership and high levels of vividness in the left posterior hippocampus. The same was true for the more distributed set of regions sensitive to body ownership during encoding. Thus, a coherent multisensory experience of a body as one’s own during encoding strengthen how core hubs of memory, including the hippocampus, reinstate the past during vivid recall. Collectively, these findings provide new insights into how the online perception of one’s own body in the center of one’s multisensory experience influences memory by binding incoming information into a common egocentric framework during encoding, which facilitates memory reinstatement.

Our full-body illusion paradigm successfully manipulated the illusory sensation of body ownership as confirmed by significant differences in illusion questionnaire ratings, threat-evoked SCR and activation of key areas associated with such illusions when comparing the synchronous and asynchronous conditions, including the premotor cortex, intraparietal cortex, lateral occipital cortex and cerebellum (Petkova et al., 2011; Guterstam et al., 2015; Preston & Ehrsson, 2016). However, we did not replicate the finding that disrupting body ownership leads to significantly impaired memory accuracy and subjective re-experiencing, as previously observed (Iriye & Ehrsson, 2022), perhaps due to a weaker illusion induction in the present study, participant fatigue from lengthened experimental sessions, and/or the distracting nature of the MR-environment. However, the lack of differences between experimental conditions in terms of memory accuracy and the subjective memory ratings clarifies the interpretation of our neuroimaging results. The neural effects of body ownership illusions on activity in areas related to encoding and reinstatement are likely due to changes in the multisensory experience of one’s own body, rather than (indirectly) mediated though changes in memory accuracy and phenomenology.

### Body Ownership Influences Memory Encoding and Reinstatement in the Hippocampus

Patterns of activity in the hippocampus distinguished between events encoded with unified compared to disrupted body ownership. The hippocampus plays a crucial role in encoding by logging distributed cortical activity elicited in response to an event, extracting associated spatial, temporal, and sensory contextual details, and merging them into a cohesive engram that can later be reinstated during retrieval (Kim, 2015; Simons, Ritchey, & Fernyhough, 2022). Our finding suggests that the information bound in the hippocampus includes multisensory signals that create a fundamental sense of body ownership, which may define a common egocentric reference frame (Ehrsson 2007 Science; (Ehrsson, 2020; Guterstam et al., 2015) that can be used to bind contextual details and centre spatial representations of an external scene (Burgess, 2006). Bergouignan and colleagues (2014) proposed that an altered sense of body ownership, self-location, and visual perspective achieved by encoding events from an out-of-body perspective impaired the ability of the hippocampus to integrate event details within this common egocentric framework, explaining the observed enhancement in hippocampal activation across repeated retrieval attempts and reduced memory vividness. However, the authors were unable to test their prediction as fMRI data was only collected during retrieval. Here, we provide evidence that body ownership, isolated from manipulations of self-location and visual perspective, is linked to memory formation in the hippocampus.

The ability of the hippocampus to categorize memories according to level of body ownership during encoding affected how the resulting memory trace was reinstated. We observed that unified body ownership and high vividness of recall increased memory reinstatement in the left posterior hippocampus. We expected body ownership to interact with vividness in the present study based on previous evidence that vividness is correlated with hippocampal activity for events encoded with atypical bodily self-awareness (Bergouignan et al., 2014), and the vividness of retrieval generally (Geib, Stanley, Wing, Laurienti, & Cabeza, 2017; Thakral, Benoit, & Schacter, 2017; Thakral, Madore, & Schacter, 2020). Thus, a coherent sense of one’s body and the vividness of recollection together enhance how the hippocampus represents the past. Memory strength is determined by the degree to which a retrieval cue matches an encoding context (*for review* see Roediger, Gallo, & Gerraci, 2002; Rugg et al., 2015). A strong match between encoding and retrieval contexts increases the chance that the index of cortical activity present at the time the remembered event was experienced will be activated in the hippocampus during retrieval, setting off a cascade of cortical activity that reinstates those original patterns and enabling the memory to be re-experienced (e.g. Hebscher, Kragel, Kahnt, & Voss, 2021). Accordingly, reinstatement was higher for events encoded with a stronger sense of body ownership and increasing levels of memory vividness both in the hippocampus, which we hypothesize initiated reactivation of the original memory, and brain regions sensitive to illusory body ownership during initial encoding, where the memory trace of patterns of activity present during encoding was stored. Hence, feelings of body ownership during encoding are likely a fundamental contextual memory cue based on spatial relationships that delineate oneself from the external world. During retrieval, feelings of a more coherent sense of body ownership consistent with those experienced at encoding in the illusion condition when the mannequin felt like oneself may facilitate access to a memory representation located in the hippocampus by increasing reinstatement. Further evidence that body ownership provides a crucial context for memory comes from our finding that patterns of activity in the parahippocampal gyrus discriminated between events experienced with stronger or weaker illusory body ownership during encoding. The parahippocampal gyrus represents information about contexts in memory, which are bound with item-level information in the hippocampus (Diana et al., 2007; Eichenbaum et al., 2007). Future studies that manipulate body ownership at both encoding and retrieval are needed to directly determine whether a coherent sense of body ownership constitutes a fundamental context for memory processing or if disrupted body ownership during encoding permanently impairs the encoding process as theorized by Bergouignan and collegues (2014).

### Patterns of Activity in the Angular Gyrus, Dorsolateral Prefrontal Cortex, and Occipital Cortex, Reflect Integration of Body Ownership into Memory

Additional brain regions outside the medial temporal lobes discriminated stronger or weaker illusory body ownership as participants formed and reinstated memories. Regions included the angular gyrus, dorsolateral prefrontal cortex, and occipital regions which have not been previously implicated in body ownership illusions and may rather reflect the incorporation of information related to own-body perception into memory encoding and reinstatement. Previous research has found that successful recall of a stimulus was predicted by similarity in patterns of brain activity in each of these regions at encoding (Hasinski & Sederberg, 2016; Kuhl et al., 2012; Lu et al., 2015; Ward et al., 2013; Xue et al., 2010, 2013). Increased similarity in patterns of activity across repetitions in prefrontal, ventral posterior parietal and sensory cortices optimizes memory by providing a more reliable, less noisy representation of encoded stimuli, which is fed to medial temporal lobe regions that rely on pattern separation mechanisms to distinguish between memories (Xue et al., 2018). We posit that the ability of frontoparietal and visual cortices to separate patterns of activity underlying memory encoding with and without body ownership may improve the distinctiveness of information sent to medial temporal lobe regions, which would then be registered as uniquely identifiable memory traces stored in the hippocampus, ultimately facilitating memory reinstatement. We suggest that the ability of the angular gyrus to discriminate between encoding of events with and without body ownership represents the integration of self-relevant bodily information into multisensory memory. Previous research has implicated this region in successful memory encoding (Lee et al., 2017; Rugg et al., 2015; Rugg & King, 2018; Uncapher & Wagner, 2009), especially for multisensory memories (Bonnici et al., 2016; Jablonowski & Rose, 2022. The angular gyrus identifies cross-modal information perceived as behaviorally self-relevant (Cabeza, Ciaramelli, Olson, & Moscovitch, 2008; Singh-Curry & Hussain, 2009; Uncapher, Benjamin Hutchinson, & Wagner, 2011), and integrates it into a common egocentric framework to support re-experiencing during retrieval (Bonnici et al., 2018; Bréchet et al., 2018; Humphreys et al., 2021; Yazar et al., 2014). We add to this literature by showing that multisensory signals underpinning a stable sense of body ownership frames how the angular gyrus encodes an event. In sum, strong compared to weak body ownership during encoding led to separate patterns of activity in parietal, frontal, and occipital areas, which may have enhanced the uniqueness of memory representations later processed in medial temporal lobe regions.

### Summary

We sought to identify brain regions where the fundamental feeling of body ownership becomes integrated within memories for events during encoding and reinstatement. Regions related to sensing a body as one’s own (i.e., left premotor and left intraparietal cortex) and memory formation (i.e., bilateral hippocampus, parahippocampal gyrus, and angular gyrus) distinguished between stronger or weaker illusory body ownership during encoding. Further, body ownership at encoding and high vividness at retrieval combined to strengthen memory reinstatement in the hippocampus and regions originally implicated in the encoding of the multisensory episode. The present study provides key insights into the role the sense of body ownership plays in creating lasting memories, which support Tulving’s (1985) idea that self-consciousness and memory are intimately linked as the sense of one’s physical self is the basis for self-consciousness.

## Acknowledgments

We would like to thank Martti Mercurio and the team at NordicNeuroLab© for their indispensable help in setting up the MR compatible HMD. We also extend our gratitude to Rouslan Sitnikov and Jonathan Berrebi at the MR Center, Department of Clinical Neuroscience, Karolinska Institutet, who implemented our scanning protocols and provided us with key equipment required to run our experiment. The project was funded by the Swedish Research Countil, Göran Gustafssons Foundation and Horzon 2020 European Research Council (Advanced Grant SELF-UNITY #787386).

## Footnotes

1 Restricting the analysis to only those participants included in the fMRI analyses (*N* = 24) led to the same pattern of results (*t*(23) = .028, Cohen’s d = .48).

2 Restricting the analysis to only those participants included in the fMRI analyses (*N* = 22) led to a marginally significant difference between the synchronous (*M* = .29, *SD* = .13) and asynchronous (*M* = .20, *SD* = .14) visuotactile stimulation conditions, *t*(21) = 1.93, *p* = 0.068, Cohen’s *d* = . 41.

3 Restricting the analysis to only those participants included in the fMRI analyses (*N* = 23) led to a main effect of coordinate, *F*(1,22) = 5.85, *p* = 0.24, η_p_^2^ = .21, revealing a greater value for mean x compared to y coordinates, *p* = .024. However, there was no significant main effect of type of visuotactile stimulation or interaction between coordinate and type of visuotactile stimulation, *p*’s > .17.

4 Restricting the analysis to only those participants included in the fMRI analyses (*N* = 24) led to the same pattern of results (*t*(23) = 1.33, *p* = .20).

5 Restricting the analysis to only those participants included in the fMRI analyses (*N* = 24) led to the same pattern of results, with main effects of session (*F*(1,22) = 24.73, *p* < .001, η_p_^2^ = .53), and detail type (*F*(1,22) = 30.84, *p* < .001, η_p_^2^ = .58), and an interaction between session and detail type (*F*(1,22) = 11.25 *p* < .003, η_p_^2^ = .34).

6 Restricting the analysis to only those participants included in the fMRI analyses (*N* = 23) led to the same interaction of session and type of visuotactile stimulation *F*(1,21) = 4.89, *p* = 0.38, η_p_^2^ = .19 without significant effects in follow-up t-tests with Bonferroni corrections, *p*’s > .48.

6 This effect was not significant when only considering data from participants included in the fMRI analyses, *F*(21) = 1.49, p = .08, η_p_^2^ = .14. No other significant main effects or interactions were observed, *p*’s > .08.

7 The main effect of testing session was also present for participants included in the fMRI analyses, *F*(1,21) = 9.87, *p* = .005, η_p_^2^=.32. There was no main effect of type of visuotactile stimulation or an interaction effect, *p’*s > .20.

8 No significant main effects or interactions were observed when only including participants in the fMRI analyses, *p*’s > .19.

