## Supplemental Table 1, Supplemental Table 2, Supplemental Table 3, Supplemental Table 4 for "The Sense of Body Ownership and the Neural Processes of Memory Encoding and Reinstatement"

### ***Supplemental Material***

Supplemental Table 1.
*Average Complexity Scores Per Sub-Category According to Group Assignment*

| Video Title | Visual | Auditory | Narrative | Emotional | Group |
|---|---|---|---|---|---|
| Moderna Museet | 2.00 | 2.50 | 1.50 | 1.36 | 1 |
| Fotografiska | 1.83 | 1.00 | 2.00 | 1.93 | 1 |
| Storkyrkan | 2.83 | 3.00 | 2.00 | 1.00 | 1 |
| Solna Centrum | 2.50 | 1.50 | 2.00 | 1.29 | 1 |
| Vasa Museet | 2.67 | 2.50 | 2.00 | 1.21 | 1 |
| Gamla Stan | 3.50 | 3.00 | 2.50 | 1.64 | 1 |
| Kungtradgården | 2.00 | 2.00 | 2.50 | 1.00 | 1 |
| Stadshuset | 2.33 | 3.00 | 2.50 | 1.14 | 1 |
| Hagaparken | 2.67 | 1.00 | 2.50 | 1.14 | 1 |
| Odenplan | 2.83 | 4.00 | 3.00 | 1.07 | 1 |
| Karlaplan | 2.33 | 1.50 | 3.00 | 1.07 | 1 |
| FilmstadenSergel | 2.67 | 3.00 | 3.50 | 1.21 | 1 |
| Central Station | 3.00 | 2.00 | 1.00 | 1.93 | 2 |
| KI Sjukhuset | 4.50 | 2.50 | 1.50 | 2.43 | 2 |
| Humlegården | 1.67 | 1.50 | 2.00 | 1.07 | 2 |
| Royal Palace | 2.17 | 3.00 | 2.00 | 1.00 | 2 |
| Strömkajen | 2.50 | 3.00 | 2.00 | 1.29 | 2 |
| Skansen | 2.33 | 1.50 | 2.50 | 1.14 | 2 |
| Stadsbiblioteket | 1.50 | 1.00 | 2.50 | 1.43 | 2 |
| Skinnarviksberget | 1.83 | 3.50 | 2.50 | 1.21 | 2 |
| Nytorget | 2.33 | 3.00 | 2.50 | 1.07 | 2 |
| Vasaparken | 2.33 | 2.50 | 2.50 | 1.00 | 2 |
| Globen | 3.00 | 3.00 | 3.50 | 1.14 | 2 |
| Grona Lund | 2.83 | 3.00 | 3.50 | 1.14 | 2 |

Supplemental Table 2.
*Differences in Average Complexity Between Groups of Video Stimuli*

| | Complexity Category | | | |
|---|---|---|---|---|
| | Visual | Auditory | Narrative | Emotional |
| Group 1 | 2.51 | 2.33 | 2.42 | 1.26 |
| Group 2 | 2.00 | 2.46 | 2.33 | 1.32 |
| *Difference* | 0.01 | -0.13 | 0.08 | -0.07 |

Supplemental Table 3.
*Average Intraclass Correlation Coeffiecients of Complexity Sub-Categories*

| Visual Complexity Measure | Intraclass Correlation Coefficient |
|---|---|
| Background | 0.80 |
| Movement | 0.73 |
| Number of Characters | 0.82 |
| Auditory Complexity Measure | |
| Background Audio | 0.88 |
| Narrative Complexity Measure | |
| Storyline | 0.62 |
| Emotional Complexity | |
| Excitement | 0.81 |
| Joy | 0.77 |
| Sadness | not enough variance |
| Anger | not enough variance |
| Disgust | not enough variance |
| Fear | not enough variance |
| Shame | not enough variance |

Supplemental Table 4.Clusters activated by memory encoding (*N* = 24)

| Anatomical region | BA | $k_E$ | $p_{FDR\text{-}corr}$ | TFCE Score | X | Y | Z |
|---|---|---|---|---|---|---|---|
| Superior Temporal Gyrus | 22 | 39776 | < 0.001 | 23232.58 | -58 | -18 | 0 |
| Superior Temporal Gyrus | 22 | | < 0.001 | 20003.64 | -56 | -10 | -6 |
| Middle Temporal Gyrus* | 21 | | < 0.001 | 19372.18 | -58 | -30 | 0 |
| Supplementary Motor Area | 6 | 167 | 0.004 | 297.57 | 2 | 12 | 62 |
| Insula | 13 | 1 | 0.034 | 327.79 | -26 | 20 | -12 |
| Dorsal Posterior Cingulate Cortex | 31 | 225 | 0.025 | 0.008 | 0 | -58 | 34 |
| Postcentral Gyrus | 1 | 1 | 0.024 | 239.54 | 48 | -20 | 54 |
| Superior Parietal Gyrus | 7 | 65 | 0.026 | 183.80 | 28 | -50 | 56 |
| Inferior Frontal Gyrus (Pars Opercularis) | 8 | 1 | 0.048 | 167.94 | 48 | 18 | 36 |
| Cerebellum Lobule IV/V | - | 1 | 0.049 | 239.54 | -20 | -32 | -30 |

* cluster extends to the hippocampus

Supplemental Table 5. Clusters activated by memory retrieval for all participants ($N$ = 29)*

| Anatomical region | BA | $k_E$ | $p_{FDR\text{-}corr}$ | TFCE Score | X | Y | Z |
|---|---|---|---|---|---|---|---|
| Premotor Cortex* | 6 | 11488 | 0.003 | 3060.19 | -26 | 8 | 52 |
| Dorsolateral Superior Frontal Gyrus | 8 | | 0.003 | 2848.31 | -16 | 18 | 40 |
| Medial Superior Frontal Gyrus | 8 | | 0.003 | 2770.88 | -8 | 20 | 44 |
| Cerebellum Lobule IX | - | 149 | 0.014 | 357.08 | 2 | -46 | -52 |
| Medulla | - | | 0.031 | 287.78 | 2 | -36 | -52 |
| Cerebellum Lobule VIII | - | 97 | 0.017 | 341.96 | 26 | -74 | -52 |
| Cerebellum Lobule VII | - | | 0.025 | 317.70 | 36 | -70 | -50 |
| Cerebellum Lobule Crus II | - | | 0.032 | 309.78 | 42 | -70 | -42 |
| Caudate | - | 54 | 0.037 | 307.14 | 2 | -2 | 22 |
| Caudate | - | | 0.045 | 294.45 | -4 | -8 | 24 |

* analysis restricted to initial 10s of each memory retrieval trial

**cluster extends to regions including dorsolateral prefrontal cortex, angular gyrus, inferior temporal cortex, and the precuneus
